# Separately prestored proteasome components to prevent polyspermy

**DOI:** 10.1101/2025.09.01.673592

**Authors:** Liying Wang, Chao Liu, Xing Wang, Yueshuai Guo, Qiuxing Zhou, Yinghong Chen, Huafang Wei, Xiaoming Huang, Shuai Liu, Wei Li, Chun So, Ling Sun, Renjie Jiao, Xuejiang Guo, Qiang Guo, Wei Li

## Abstract

To ensure genome stability across generations, an egg must be fertilized by only a single sperm. However, how sperm actively contribute to preventing polyspermy has remained unknown. Here, we demonstrate that testis-specific 20S proteasome core particles (CPs) are pre-stored within the sperm heads of both mice and humans. Upon fertilization, these testis-specific 20S CPs assemble with oocyte-derived 19S regulatory particles (RPs), forming chimeric proteasomes that efficiently degrade Fetuin B, which is ubiquitinated by the E3 ligase MARCH3. The clearance of Fetuin B triggers ZP2 cleavage and subsequent zona pellucida hardening, thereby blocking additional sperm entry. The newly assembled chimeric proteasome in the zygote represents a novel strategy to prevent polyspermy during fertilization in mammals.

## INTRODUCTION

During fertilization in mammals, millions of sperm compete for their destination, the egg. If multiple sperm claim the egg, polyspermy leads to early embryo arrest or abortion in mammals.^1,2^ To prevent polyspermy, both the oolemma (egg plasma membrane) and zona pellucida undergo biochemical alterations to ensure the egg can accommodate only a single sperm, thereby blocking polyspermy and preventing polyploidy.^3,4^

In mammals, polyspermy is prevented by the zona pellucida block and the membrane block. JUNO is responsible for the membrane block and is shed from the oocyte’s plasma membrane approximately 40 minutes after fertilization.^5–7^ Additionally, the release of CD9-containing vesicles contributes to the establishment of the membrane block.^8^ Upon fertilization, the oocyte’s internal calcium concentration increases, prompting cortical granules to fuse with the oolemma and exocytose their contents.^3,9^ Ovastacin proteolytically processes ZP2, destroying the sperm-binding region within ZP2 and rendering eggs refractory to further sperm adhesion and fertilization.^10,11^ Fetuin B, a cystatin superfamily protein (thiol protease inhibitor), has been shown to directly inhibit the physiological release and proteolytic action of ovastacin, thereby preventing premature and inappropriate hardening of the zona pellucida.^12–14^ Zinc exocytosed from egg cortical granules also affects sperm motility, enabling a transient block to zona pellucida penetration.^15^ Additionally, transglutaminase 2 (TGM2) crosslinks ZP3, inducing the hardening of the zona pellucida after fertilization to prevent sperm penetration and polyspermy.^16,17^ While some maternal factors that block sperm entry have been characterized, the upstream signaling cascade that triggers this response remains elusive. Given this event must be triggered by sperm penetration, some paternal factor(s) should be involved in this quick cascade.

Up to now, only a single paternal factor, PLCζ, was found to be involved in this process,^18,19^ and other paternal factors are expected to be found during fertilization. A mature sperm is a highly differentiated cell that is made up of three parts: the acrosome, the head, and the tail.^20^ The acrosome is a cap-like structure covering the anterior portion of the sperm head. During fertilization, acrosomal enzymes with chymotryptic activity are released to clear a path to the egg for the incoming sperm, resulting in the loss of the sperm’s acrosome and plasma membrane components.^21,22^ Following entry into the egg cortex, the sperm nucleus and tail are incorporated into the egg cytoplasm. The sperm nucleus contains highly condensed chromatin, where DNA is tightly packaged by protamines—small (∼50 amino acids), arginine-rich nuclear proteins that neutralize DNA charge and facilitate chromatin compaction through disulfide crosslinking.^23,24^ The decondensation of protamine-bound DNA occurs, accompanied by the initiation of protamine-to-histone exchange are detected within one hour after fertilization.^25^ The sperm tail starts to disappear at the late 8-cell stage and becomes undetectable after the 32-cell stage.^26^ In the sperm head, nuclear vacuoles appear as small clear spaces within the condensed chromatin of the sperm nucleus^27^ and may contain cytoskeletal proteins and cytoplasmic enzymes.^28^ Considering that the prevention of polyspermy is a rapid process, paternal factors might also be stored in nuclear vacuoles.

To investigate the potential function(s) of sperm in polyspermy prevention, we used cryo-electron tomography (cryo-ET) technology to visualize human and mouse sperm heads, and found that an abundance of 20S CPs were stored in the nuclear vacuoles of the sperm head. After fertilization, these 20S CPs are integrated with 19S RPs in oocytes. We show that the newly assembled chimeric proteasomes promote the degradation of Fetuin B, which is polyubiquitinated by the ubiquitin E3 ligase MARCH3, thus accelerating ZP2 cleavage, hardening the zona pellucida, and ultimately preventing polyspermy. Our study demonstrates, in both human and mouse, separately prestored 20S CP from sperm and 19S RP from oocytes assemble into chimeric proteasomes to prevent polyspermy after fertilization.

## RESULTS

### 20S CPs are stored in the sperm head

Cryo-electron tomography (cryo-ET) has revolutionized our ability to visualize cellular structures at near-native conditions, enabling three-dimensional imaging at molecular resolution.^29^ To explore the potential role of nuclear vacuoles in preventing polyspermy, we employed cryo-ET to visualize the head region of cryo-focused ion beam-thinned human sperm (Figure 1). The human sperm head is predominantly occupied by an electron-dense region (Figure 1A). In addition to highly condensed chromatin packaged with protamines, we found electron-transparent areas, which were previously identified as nuclear vacuoles.^27,28^ Upon closer inspection of tomographic cross-sections, these vacuoles are densely populated with macromolecules that exhibit a ring-like structure of approximately 10 nm in diameter (Figure 1B). To elucidate the identity of these macromolecules, we aligned and averaged the particles, revealing a distinctive barrel-shaped structure with 7-fold symmetry, characteristic of 20S CPs within the proteasome (Figure 1C).

**Figure 1.**
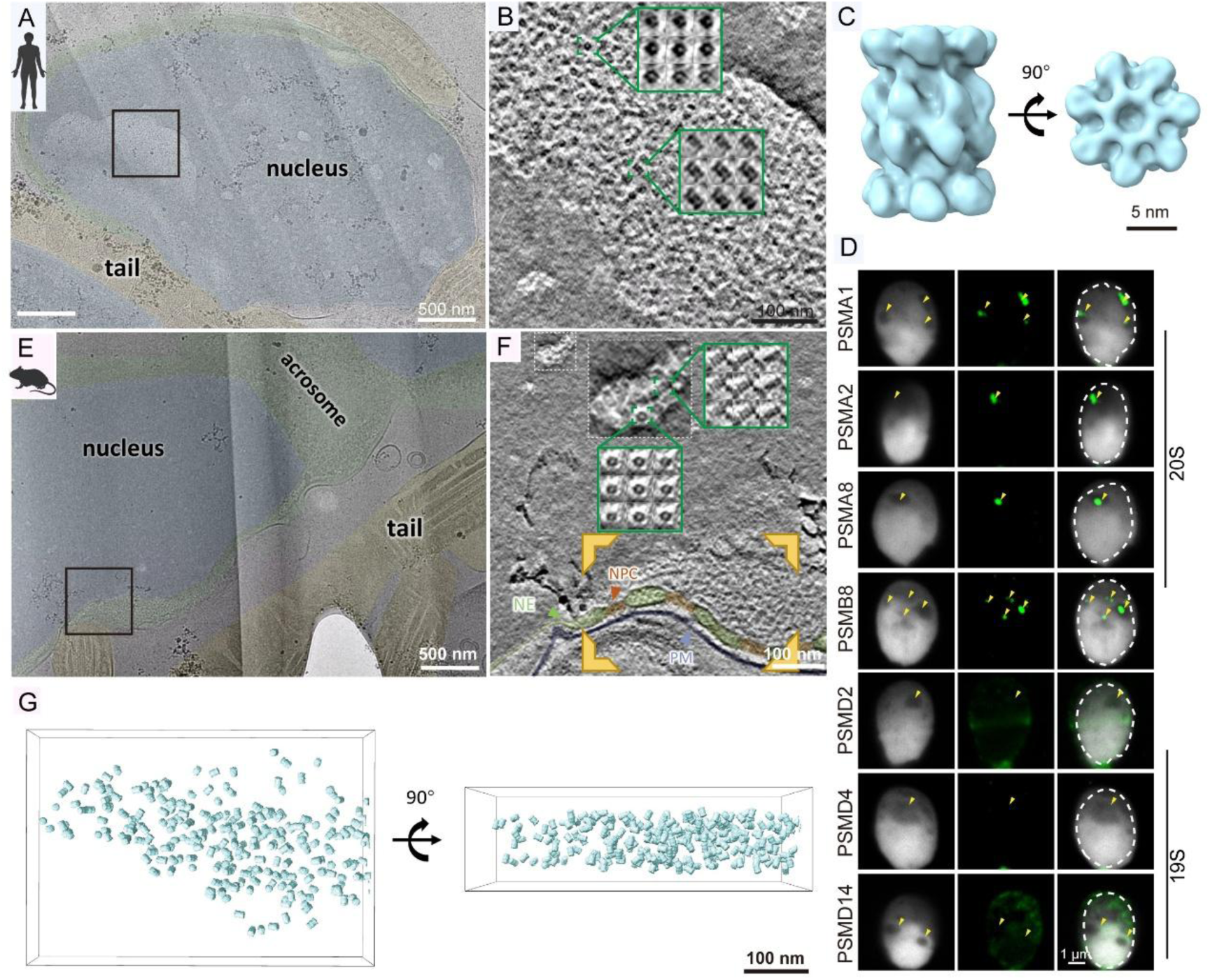
20S CPs form foci in sperm nucleus. (A) A low-magnification TEM image of a lamella showing the head of a human sperm cell. Nuclear regions are marked with transparent blue shading, and acrosome regions are marked with transparent green shading. The black box, containing a specific focus, is enlarged and shown in (B). (B) A slice from the reconstructed tomogram of the region targeted in (A). Two representative 20S CPs are displayed in green boxes, visualized as 1.58 nm Z slices within each volume. (C) Different views of the subtomogram averaging map for the human 20S CP are presented. (D) Representative images of human spermatozoa immunostained with proteasome components (green) and counterstained with DAPI. (E) A low-magnification TEM image of a mouse sperm lamella showing a sperm cell. Tail regions are highlighted in yellow. A representative region containing these foci is marked by a black box and selected for cryo-ET analysis. (F) A slice from the reconstructed tomogram of the targeted region in (E). The nuclear vacuole within the white dashed box was magnified to a 100 nm square, from which two representative 20S CPs are displayed in green boxes, shown as a 1.58 nm Z slice within each volume. NE, nuclear envelope; NPC, nuclear pore complex; PM, plasma membrane. (G) The clustered mouse 20S CPs detected within the yellow bounding frame in (F) are visualized as blue surfaces and presented in different orientations.

The 26S proteasome serves as the primary proteolytic complex responsible for regulated protein degradation in eukaryotic cells. This conserved complex is made up of two functional components: a barrel-shaped 20S CP, which harbors the catalytic activity, and a 19S RP (Figure S1A). The RPs are responsible for substrate recognition, unfolding, and translocation into the core particle for subsequent proteolysis.^30^ The 20S CP is composed of four stacked heptameric rings arranged in an (α1-α7) (β1-β7) (β1-β7) (α1-α7) configuration. The inner two catalytic β rings carry out proteolytic reactions, while the outer two structural α rings interact with regulatory particles.^31^ Ubiquitin and proteasome systems have been implicated in many reproductive processes,^32–34^ with some results hinting at proteasome functions in fertilization.^35–37^

We further used immunofluorescence staining to reveal that 20S CP components (PSMA1, PSMA2, PSMA8, and PSMB8) localized to nuclear vacuoles in human sperm (Figure 1D), consistent with our structural findings. Interestingly, 19S RP components (PSMD2, PSMD4, and PSMD14) were absent from these vacuoles. Previous structural studies have demonstrated that the proteasome primarily exists as a 26S complex in various organisms.^38^ The enrichment of 20S CPs in sperm nuclear vacuoles may suggest a novel storage mechanism.

To investigate the conservation of this phenomenon, we examined mouse sperm using cryo-ET and subtomogram averaging. Similar to human sperm, mouse sperm exhibited nuclear vacuoles containing 20S CPs, albeit smaller in size but more numerous (Figures 1E-G and Figures S1B-D). Immunofluorescence staining of mouse germ cells revealed that 20S CP components were present in the nucleus at all development stages (Figures S2A-C). In contrast, 19S RP components were detectable in spermatocytes and round spermatids but absent at later stages (sperm head elongation and differentiation; Figures S2D-E). Western blotting indicated that 20S CP components, but not 19S RP components, were stored in the sperm heads (Figure S2F). We also evaluated the localization of proteasome components in oocytes and embryos, finding that PSMA1, PSMB8, PSMD2 and PSMD4 were present in oocytes and embryos, while PSMA8 was detected in embryos but not oocytes (Figures S3A-E). These findings indicate that 20S CP may be stored during sperm maturation.

### Depletion of paternal 20S CP components leads to polyspermy

The mouse is an excellent model for studying the functional role of stored 20S CP components in the sperm head during fertilization. We subsequently generated conditional knockout alleles for genes encoding the 20S CP components PSMA1 and PSMA8 by flanking their exons with loxP sites. These two genes were specifically knocked out in germ cells by crossing floxed mice with Protamine-Cre mice, which express Cre recombinase in haploid spermatids.^39^ We hereafter refer to germ cell-specific knockout mice for *Psma1* and *Psma8* as *Prm-Psma1*^-*/*-^ and *Prm-Psma8*^-*/*-^ mice. Immunoblotting against PSMA1 and PSMA8 showed significantly reduced protein levels in mature sperm of *Prm-Psma1*^-*/*-^ and *Prm-Psma8*^-*/*-^ mice compared to control (*Psma1^Flox/Flox^*and *Psma8^Flox/Flox^*) mature sperm (Figures S4A-B). Neither testis size nor total sperm count were affected in *Prm-Psma1*^-*/*-^ and *Prm-Psma8*^-*/*-^ mice (Figures S4C-E). Hematoxylin-eosin staining revealed that the seminiferous tubules and cauda epididymis of *Prm-Psma1*^-*/*-^ and *Prm-Psma8*^-*/*-^ mice displayed normal structures, with no obvious defects compared to control mice (Figures S4F-G). Additionally, we performed single-sperm immunofluorescence using the acrosome-specific marker peanut agglutinin (PNA), with DAPI co-staining to show the sperm nucleus. Sperm morphology was characterized using confocal microscopy, revealing that *Prm-Psma1*^-*/*-^, *Prm-Psma8*^-*/*-^, and control sperm exhibited normal morphology (Figures S4H-I). These results indicate that the depletion of PSMA1 or PSMA8 is dispensable for later stages of spermatogenesis.

We next examined the potential impact of germline-specific knockout of 20S CP components on male fertility. Fertility testing showed that average litter sizes were significantly decreased in *Prm-Psma1*^-*/*-^ and *Prm-Psma8*^-*/*-^ males compared to control males (Figure 2A). Thus, males with germ cell-specific knockout of the 20S CP components PSMA1 and PSMA8 are subfertile. We subsequently performed *in vitro* fertilization (IVF) using 20S CP component conditional knockout (cKO) sperm and wild-type (WT) oocytes to generate *Psma1^♀+/♂-^* or *Psma8^♀+/♂-^* embryos, which led to the paternal depletion (♂-) but maternal presence (♀+) of PSMA1 or PSMA8 proteins. For the control group (*20S CP ^♀+/♂+^*), WT oocytes were fertilized with *Psma1^Flox/Flox^*or *Psma8^Flox/Flox^* sperm (Figure 2B). Both *Psma1^♀+/♂-^* and *Psma8^♀+/♂-^* embryos showed early developmental arrest compared to control embryos (Figures 2C-D). Conspicuously, we observed extra sperm residing in *Psma1^♀+/♂-^* and *Psma8^♀+/♂-^* embryos (Figure 2C). Immunofluorescence staining indicated that embryos from fertilization with cKO sperm showed more than three pronuclei, whereas *20S CP ^♀+/♂+^* did not (as *20S CP ^♀+/♂+^* shows two pronuclei) (Figures 2E-F). These results demonstrate that knockout of 20S CP components PSMA1 and PSMA8 in male germ cells leads to polyspermy in embryos, suggesting that prestored 20S CP components in sperm somehow function as paternal factors that prevent polyspermy.

**Figure 2.**
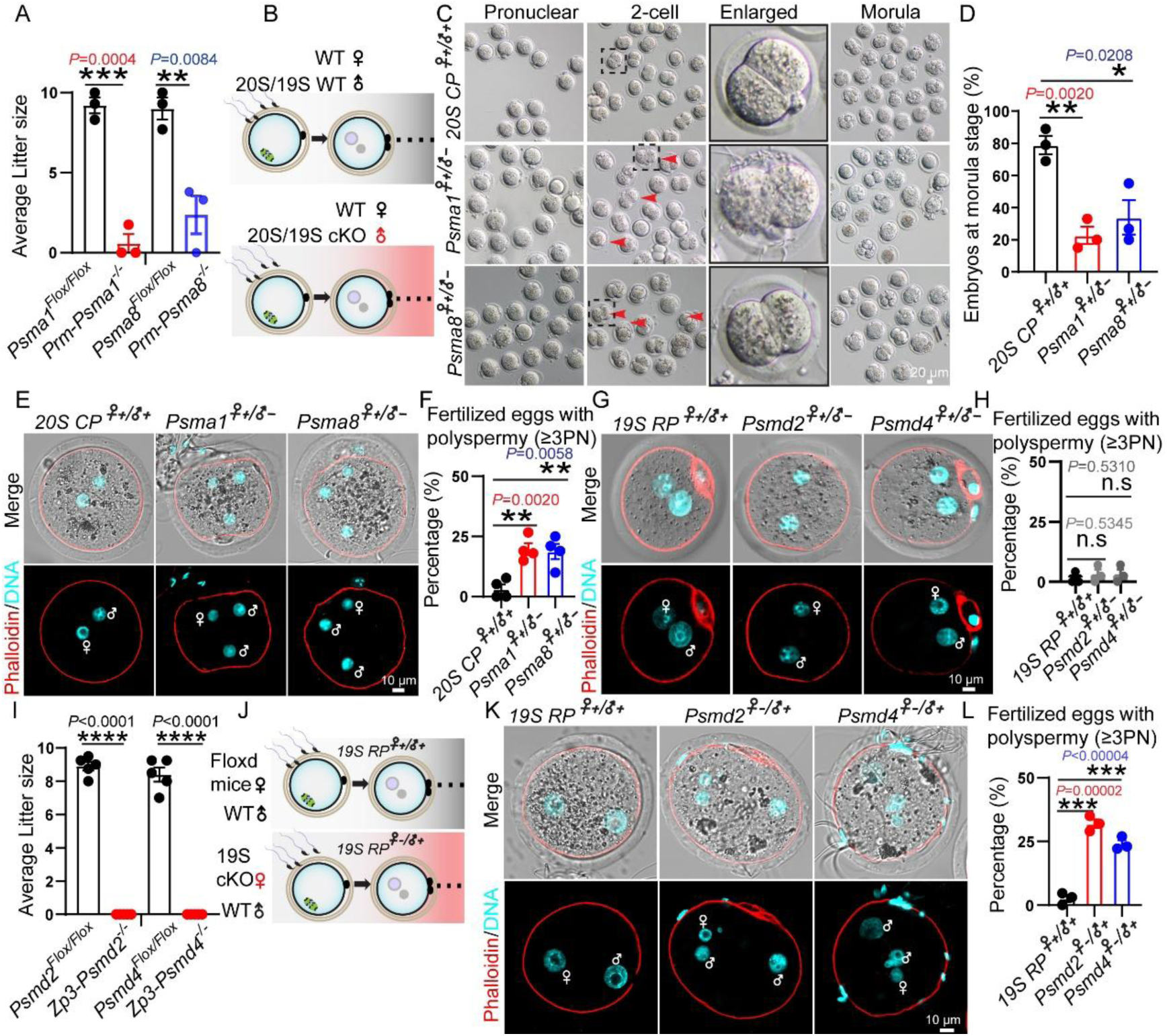
Depletion of proteasome components leads to polyspermy. (A) The average litter size of *Psma1^Flox/Flox^*, 9.20 ± 0.49 (n = 3 independent experiments); *Prm-Psma1^−/−^*, 0.58 ± 0.58(n = 3 independent experiments); *Psma8^Flox/Flox^*, 9.00 ± 0.68 (n = 3 independent experiments); *Prm-Psma8^−/−^*, 2.37 ± 1.19 (n = 3 independent experiments). Data are presented as the mean ± SEM. Two-tailed unpaired Student’s t test, ***p < 0.001, **p < 0.01. (B) Schematic of collecting samples for control and knockout mice. (C) Distribution stages of IVF embryos from pronuclear until morula. Arrows indicate abnormal embryos. (D) Percentage of embryos at the morula stage from control, paternal *Psma1*-depleted and paternal *Psma8*-depleted groups. *20S CP^♀+/♂+^*, 79.00% ± 5.77% (n = 3 independent experiments, total oocytes = 61); *Psma1^♀+/♂-^*, 22.67% ± 5.36% (n = 3 independent experiments, total oocytes = 97); *Psma8^♀+/♂-^*, 33.90% ± 10.73% (n = 3 independent experiments, total oocytes = 50). Data are presented as the mean ± SEM. Two-tailed unpaired Student’s t test, **p < 0.01, *p < 0.05. (E) Representative images of pronuclei stained with Phalloidin (red) and DAPI (cyan) in control and paternal *Psma1*-depleted and paternal *Psma8*-depleted groups after 6 hours post IVF. Female and male symbols indicate the female and male pronuclei, respectively. (F) Percentage of fertilized oocytes with polyspermy (≥3PN) was quantified in **e**. *20S CP^♀+/♂+^*, 3.23% ± 1.93% (n = 4 independent experiments, total oocytes = 82); *Psma1^♀+/♂-^*, 19.68% ± 2.49% (n = 4 independent experiments, total oocytes = 79); *Psma8^♀+/♂-^*, 18.75% ± 3.17% (n = 4 independent experiments, total oocytes = 85). Data are presented as the mean ± SEM. Two-tailed unpaired Student’s t test, **p < 0.01. (G) Representative images of pronuclei stained with Phalloidin (red) and DAPI (cyan) in *19S RP^♀+/♂+^*, *Psmd2^♀+/♂-^* and *Psmd4^♀+/♂-^*groups after 6 hours post IVF. (H) Percentage of fertilized oocytes with polyspermy (≥3PN) was quantified in (G). *19S RP^♀+/♂+^*, 1.43% ± 1.43% (n = 3 independent experiments, total oocytes = 63); *Psmd2^♀+/♂-^*, 3.13% ± 2.05% (n = 3 independent experiments, total oocytes = 91); *Psmd4^♀+/♂-^*, 3.17% ± 2.09% (n = 3 independent experiments, total oocytes = 98). Data are presented as the mean ± SEM. Two-tailed unpaired Student’s t test, n.s., non-significant. (I) The average litter size of *Psmd2^Flox/Flox^*, 8.90 ± 0.26 (n = 5 independent experiments); *Zp3-Psmd2^−/−^*, 0.00 ± 0.00 (n = 5 independent experiments); *Psmd4^Flox/Flox^*, 8.40 ± 0.42 (n = 5 independent experiments); *Zp3-Psmd4^−/−^*, 0.00 ± 0.00 (n = 5 independent experiments). Data are presented as the mean ± SEM. Two-tailed unpaired Student’s t test, ****p < 0.0001. (J) Schematic of collecting samples in control and knockout mice. (K) Representative images of pronuclei stained with Phalloidin (red) and DAPI (cyan) in control, maternal *Psmd2*-depleted and maternal *Psmd4*-depleted mice after 6 hours post IVF. Female and male symbols indicate the female and male pronuclei, respectively. (L) Percentage of fertilized oocytes with extra sperm in the perivitelline space was quantified in (K). *19S RP^♀+/♂+^*, 2.60% ± 1.38% (n = 3 independent experiments, total oocytes = 84); *Psmd2^♀-/♂+^*, 32.00% ± 1.73% (n = 3 independent experiments, total oocytes = 71); *Psmd4^♀-/♂+^*, 24.07% ± 1.49% (n = 3 independent experiments, total oocytes = 88).

Notably, knocking out 19S RP components PSMD2 and PSMD4 in spermatids had no significant effect on testis size, total sperm count, seminiferous tubule and cauda epididymis structure, or sperm morphology (Figures S4J-P). Fertility testing showed that *Prm-Psmd2*^-*/*-^ and *Prm-Psmd4*^-*/*-^ male mice displayed no signs of subfertility (Figure S4Q). Additionally, PSMD2-or PSMD4-depleted sperm achieved normal fertilization, and the percentage of polyspermy in embryos remained unchanged (Figures 2G-H). These results suggest that 20S CP components, but not 19S RP components, from sperm are essential for fertilization.

### Depletion of maternal 19S RP components leads to polyspermy

To explore the potential role(s) of maternal proteasome components in preventing polyspermy, we examined the functional impact of selectively depleting PSMA1, PSMD2, or PSMD4 from oocytes. We crossed *Psma1* flox, *Psmd2* flox, or *Psmd4* flox mice with *Zp3-Cre* transgenic mice to generate *Psma1^Flox/Flox^*; *Zp3-Cre*, *Psmd2^Flox/Flox^*; *Zp3-Cre*, and *Psmd4^Flox/Flox^*; *Zp3-Cre* mice, hereafter referred to as *Zp3-Psma1*^-*/*-^, *Zp3-Psmd2*^-*/*-^, and *Zp3-Psmd4*^-*/*-^ mice. *Zp3-Cre* expression begins in oocytes at the primary follicle stage, starting around postnatal day 5 (PN5) .^40^ Immunoblotting against PSMA1, PSMD2, and PSMD4 showed significantly reduced protein levels in germinal vesicle (GV) oocytes of the respective conditional knockout compared to control oocytes (*Psma1^Flox/Flox^, Psma8^Flox/Flox^, Psmd2^Flox/Flox^*, and *Psmd4^Flox/Flox^*) (Figure S5A-F). Fertility testing showed that female mice lacking PSMA1, PSMD2, or PSMD4 were completely infertile (Figure 2I, Figure S5G). Moreover, the size of *Zp3-Psma1*^-*/*-^ ovaries was smaller than that of the controls, and no oocytes were released from ovaries upon superovulation (Figures S5H-I), suggesting that PSMA1 depletion in ovaries may result in premature ovarian insufficiency.

PSMD2 depletion had little effect on ovarian size at 4 weeks, but ovaries became smaller at 8 weeks (Figure S5J). PSMD4 depletion had no obvious effect on ovarian size (Figure S5K). We then combined *Zp3-Psmd2*^-*/*-^ or *Zp3-Psmd4*^-*/*-^ oocytes with WT sperm through IVF to generate *Psmd2^♀-/♂+^* or *Psmd4^♀-/♂+^* embryos (Figure 2J). Immunofluorescence staining showed more than three pronuclei in the *Psmd2^♀-/♂+^* embryos (Figure 2K). Similar results were observed with *Zp3-Psmd4*^-*/*-^ female mice, where extra sperm resided in the perivitelline space of *Psmd4^♀-/♂+^*embryos after 6 hours and 24 hours post IVF (Figures S5L-M). Immunofluorescence staining showed more than three pronuclei in the *Psmd4^♀-/♂+^*embryos (Figures 2K-L). Additionally, after mating WT and *Zp3-Psmd4*^-*/*-^ female mice with WT males, E0.5 *Psmd4^♀-/♂+^* embryos flushed from the oviducts showed extra sperm in the perivitelline space (Figure S5N). Embryos from fertilization with 19S RP cKO oocytes showed more than three pronuclei, whereas control group did not (as control group shows two pronuclei) (Figures S5O-P). These results demonstrate that the depletion of maternal 19S RP components PSMD2 and PSMD4 results in polyspermy.

### 20S CP from sperm and 19S RP from oocytes assemble into chimeric proteasomes after fertilization

Given our findings that polyspermy results from either loss of paternal 20S CP components or loss of maternal 19S RP components, and since the sperm nucleus stores 20S CP but not 19S RP, we propose that sperm and oocytes separately store 20S CP and 19S RP. These separated components assemble into chimeric proteasomes upon fertilization to prevent polyspermy.

We examined this idea by generating knock-in mice bearing proteasome subunits labeled with eGFP or mCherry fluorescent tags. These subunits included 20S CP components PSMA1 and PSMA8 and 19S RP components PSMD2 and PSMD4. We first demonstrated PSMA1-eGFP, PSMD2-mCherry, PSMA8-mCherry, and PSMD4-eGFP proteins were expressed in testis in heterozygous knock-in mice (Figures S6A-D). Consistent with cryo-ET and immunofluorescence results, 20S CP components PSMA1 and PSMA8 were stored in the nucleus of spermatozoa, whereas 19S RP components PSMD2 and PSMD4 were not (Figure 3A), indicating fluorescent labeling does not affect 20S CP storage in the sperm head. PSMA1, PSMD2, and PSMD4 were all detected in metaphase II (MII) stage oocytes (Figure 3B). To our surprise, we observed that PSMD4-eGFP rapidly formed foci surrounding PSMA8-mCherry in sperm and formed chimeric proteasomes during fertilization. Notably, the newly assembled chimeric proteasomes were distributed to oocytes approximately 30-40 minutes after fertilization (Figures 3C-D; Supplementary Video 1). We also observed that PSMD2-mCherry rapidly formed foci surrounding PSMA1-eGFP in sperm and formed chimeric proteasomes during fertilization (Figures S6E-F). Stimulated emission depletion microscopy (STED) further confirmed the formation of chimeric proteasomes in fertilized oocytes (Figures 3E-G). Co-immunoprecipitation analysis revealed that the testis-specific proteasome 20S CP subunit PSMA8 or PSMA7 (paralog of PSMA8) physically interacts with 20S CP components PSMA1, PSMA2, and PSMB8, as well as 19S RP components PSMD2, PSMD4, and PSMD14 (Figure S6G). The proximity ligation assay (PLA) also demonstrated an interaction between PSMA1 and PSMD2 in embryos after fertilization of wild-type oocytes with PSMA1-eGFP sperm (Figure 3H). Together, these results suggest that testis-specific 20S CPs are integrated with oocyte-derived 19S RPs to form chimeric proteasomes that are then distributed to the oocytes, potentially preventing polyspermy during fertilization.

**Figure 3.**
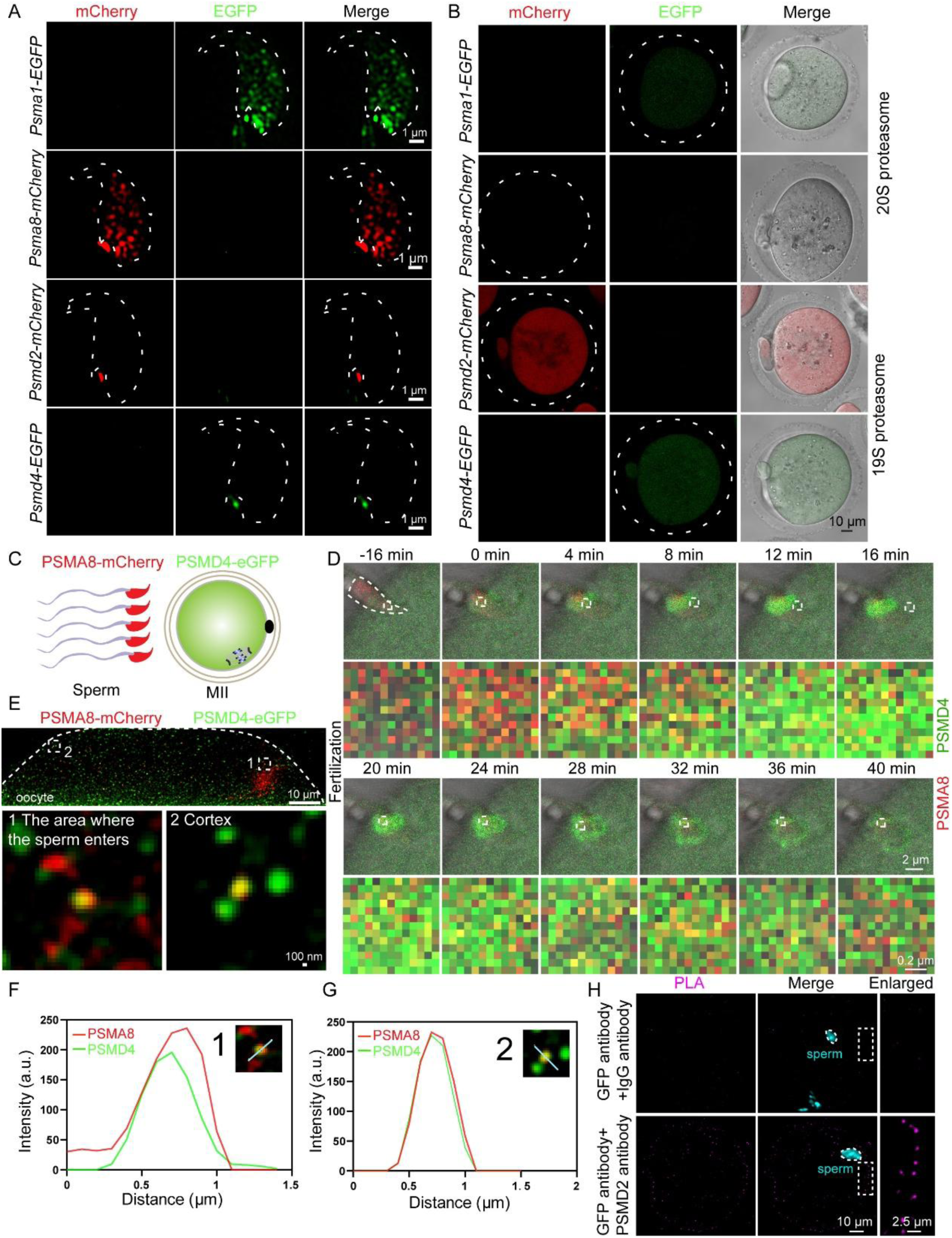
Assembly of chimeric proteasomes during fertilization. (A) Super-resolution microscopy images show the localization of proteasome components in the spermatozoa of PSMA1-eGFP, PSMA8-mCherry, PSMD2-mCherry, and PSMD4-eGFP knock-in mice. Dotted lines indicate the sperm heads. (B) Representative images display the localization of proteasome components in MII stage oocytes of PSMA1-eGFP, PSMA8-mCherry, PSMD2-mCherry, and PSMD4-eGFP knock-in mice. (C) Schematic of *in vitro* fertilization process by PSMA8-mCherry and PSMD4-eGFP knock-in mice. (D) Live imaging results showed that PSMD4-eGFP rapidly formed foci surrounding PSMA8-mCherry sperm and led to the formation of chimeric proteasomes during fertilization. (E) Representative STED microscope images display the newly assembled proteasomes during fertilization. (F) and (G) Line-scan analysis (cyan line) was performed using Image J software in (E). (H) Proximity ligation assay detection of PSMA1-PSMD2 interaction in embryos. Wild-type (WT) oocytes were fertilized with PSMA1-eGFP sperm for 1.5 h, followed by PLA to assess protein-protein interaction.

### Depletion of proteasome components results in Fetuin B accumulation and uncleaved ZP2 in embryos

Following fertilization, the cleavage of ZP2 leads to the hardening of the zona pellucida, preventing further sperm from binding to the oocyte.^10,13^ To investigate potential biochemical mechanisms through which proteasome components contribute to preventing polyspermy, we first assessed zona pellucida hardening by measuring chymotrypsin-mediated zona pellucida digestion time in WT, *Psma8^♀+/♂-^*, and *Psmd4^♀-/♂+^* two-cell embryos. The zona pellucida digestion time of *Psma8^♀+/♂-^* and *Psmd4^♀-/♂+^* two-cell embryos was significantly shorter than that of WT two-cell embryos, suggesting an absence of zona pellucida hardening in *Psma8^♀+/♂-^* and *Psmd4^♀-/♂+^*embryos (Figures S7A-B).

After fertilization, the ZP2 precursor was initially reported to be cleaved into a form that retains disulfide bonds and was a determinant of sperm binding to the surface of the zona pellucida.^10^ With our control group of two-cell embryos, ZP2 was cleaved and served as a positive control. In contrast, immunoblotting of *Psma8^♀+/♂-^*, *Psmd2^♀-/♂+^*, and *Psmd4^♀-/♂+^* two-cell embryos showed that ZP2 had an uncleaved mass of 120 kDa (Figures 4A-B, Figure S7C). We subsequently performed a sperm binding assay and found that capacitated mouse sperm bound to *Psma1^♀+/♂-^* MII oocytes (13.95 ± 5.20 SEM sperm/egg), *Psma8^♀+/♂-^* MII oocytes (33.29 ± 7.91 SEM sperm/egg), and *Psmd4^♀-/♂+^* MII oocytes (132.40 ± 9.08 SEM sperm/egg), but did not bind to WT control group of MII oocytes with one hour after fertilization (2.35 ± 0.66 SEM sperm/egg) (Figures 4C-F, Figures S7D-E). Together, these results demonstrate that depletion of paternal 20S CP components or maternal 19S RP components leads to uncleaved ZP2, an absence of zona pellucida hardening, and ultimately to polyspermic embryos.

**Figure 4.**
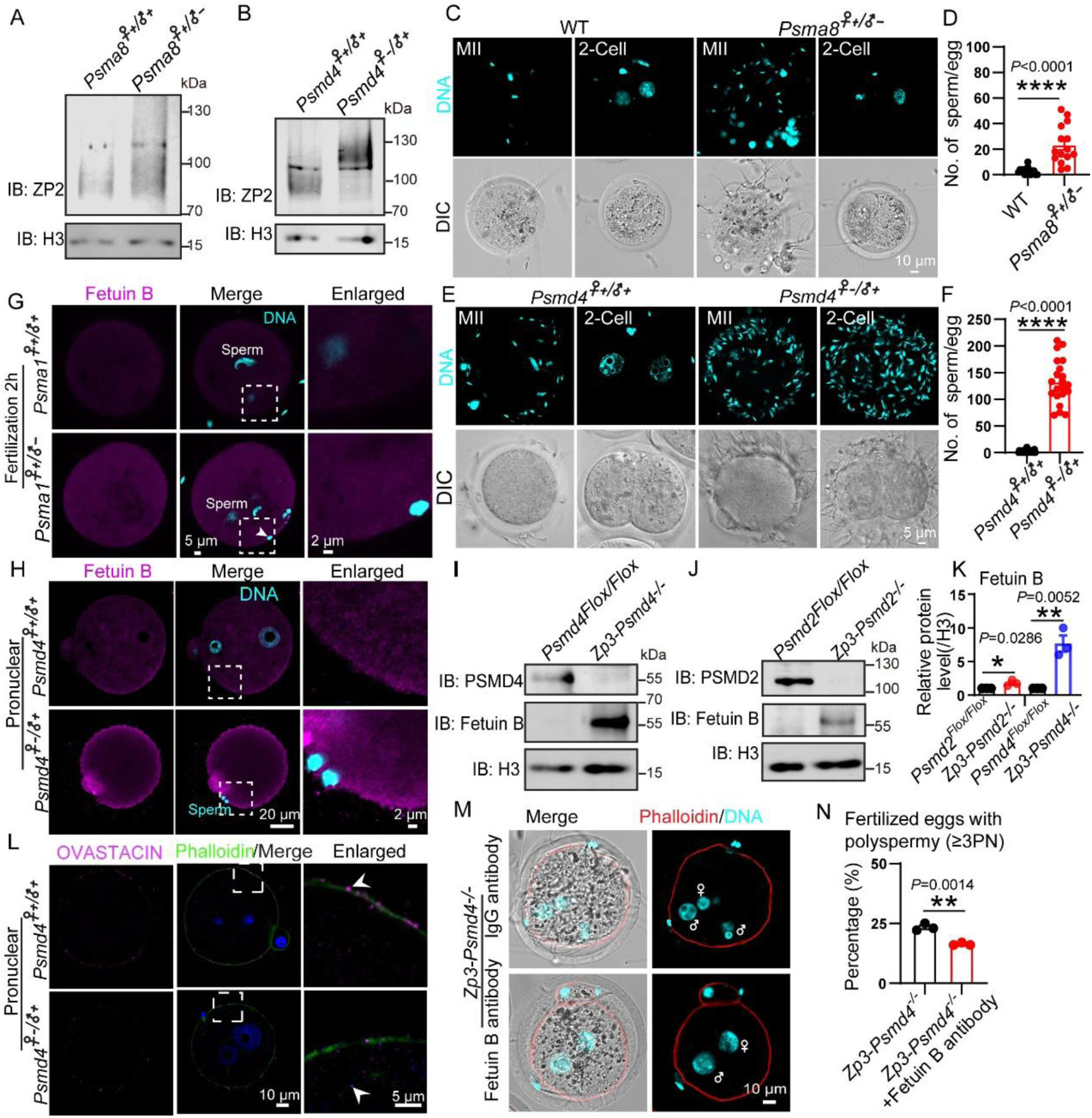
Depletion of proteasome components result in the accumulation of Fetuin B and uncleaved ZP2 in embryos. (A and B) ZP2 remains uncleaved in maternal PSMD4-depleted embryos or paternal PSMA8-depleted embryos after *in vitro* fertilization. Western blots were probed with anti-ZP2 and anti-H3 antibodies. H3 served as a loading control. (C) MII oocytes and two-cell embryos from WT mice were incubated with both *Psma8^Flox/Flox^* and PSMA8-depleted capacitated sperm for 1 hour. (D) The number of *Psma8^Flox/Flox^* and PSMA8-depleted sperm binding to the surface of the zona pellucida. WT, 2.67 ± 0.71 (n = 15); *Psma8^♀+/♂-^*, 23.27 ± 3.90 (n = 15). Data are presented as the mean ± SEM. Two-tailed unpaired Student’s t test, ****p < 0.0001. (E) Representative images of sperm binding to the zona pellucida of PSMD4-depleted oocytes and embryos. MII oocytes and two-cell embryos from control and PSMD4-depleted mice were incubated with capacitated sperm for 1 hour. The oocytes and embryos were then stained with DAPI (cyan). (F) The number of sperm binding to the surface of ZP surrounding oocytes in (E). *Psmd4^♀+/♂+^*, 2.00 ± 0.56 (n = 21); *Psmd4^♀+/♂-^*, 132.40 ± 9.08 (n = 21). Data are presented as the mean ± SEM. Two-tailed unpaired Student’s t test, ****p < 0.0001. (G) Representative images showing Fetuin B localization in control and paternal PSMA1-depleted embryos after *in vitro* fertilization for 2 hours. Arrows indicate extra sperm inside the plasma membrane. (H) Representative images showing Fetuin B localization in control and maternal PSMD4-depleted embryos during the pronuclear stage. (I) Fetuin B protein levels in *Zp3-Psmd4^−/−^* oocytes were significantly increased compared to control oocytes. Western blots were probed with anti-PSMD4, anti-Fetuin B and anti-H3 antibodies. H3 served as a loading control. (J) Fetuin B protein levels in *Zp3-Psmd2^−/−^* oocytes were significantly increased compared to control oocytes. Western blots were probed with anti-PSMD2, anti-Fetuin B and anti-H3 antibodies. H3 served as a loading control. (K) Relative protein levels of Fetuin B in *Psmd2^Flox/Flox^*and *Zp3-Psmd2^−/−^* oocytes, *Psmd4^Flox/Flox^* and *Zp3-Psmd4^−/−^* oocytes. (n = 3 independent experiments). Data are presented as the mean ± SEM. Two-tailed unpaired Student’s t test, *p < 0.05, **p < 0.01. (L) Representative images showing ovastacin localization in control and maternal PSMD4-depleted embryos during the pronuclear stage. Arrows indicate ovastacin signal in the perivitelline space or cytoplasm. (M) Representative images of embryos in *Psmd4*-depleted mice after 6 hours post IVF. Fetuin B antibody was used to block Fetuin B during fertilization. Female and male symbols indicate the female and male pronuclei, respectively. (N) Percentage of fertilized oocytes with polyspermy (≥3PN) was quantified in M. *Zp3-Psmd4^−/−^*, 23.40% ± 0.83% (n = 3 independent experiments, total oocytes =72); *Zp3-Psmd4^−/−^+*Fetuin B antibody, 16.33% ± 0.33% (n = 3 independent experiments, total oocytes =79). Data are presented as the mean ± SEM. Two-tailed unpaired Student’s t test, **p < 0.01.

Fetuin B is a plasma protein that directly inhibits ovastacin activity and is essential for preventing premature hardening of the zona pellucida in oocytes.^12^ Immunofluorescence staining and immunoblotting showed Fetuin B accumulated in embryos upon depletion of paternal 20S CP components PSMA1 or PSMA8 (Figure 4G, Figures S7F-G). The depletion of maternal 19S RP components PSMD2 or PSMD4 also led to the accumulation of Fetuin B and defects in ovastacin release (Figures 4H-L). Additionally, functional assays revealed that blocking Fetuin B by adding a Fetuin B antibody during fertilization partially rescued the polyspermy phenotype in PSMD4-depleted mice (Figure 4M). The percentage of fertilized oocytes with polyspermy (≥3PN) was significantly decreased in *Zp3-Psmd4^−/−^* mice compared to control mice (Figure 4N). Together, these results demonstrate that the depletion of paternal 20S CP components or maternal 19S RP components leads to Fetuin B accumulation, which results in uncleaved ZP2 and polyspermy in embryos.

### The E3 ligase MARCH3 mediates polyubiquitination of Fetuin B

The observation that depletion of proteasome components leads to Fetuin B accumulation suggests that Fetuin B may be degraded by the proteasome. First, we performed cycloheximide (CHX) chase assays in HEK293T and found that adding the proteasome inhibitor MG132 resulted in strong accumulation of Fetuin B compared to the control group (Figures 5A-B). Moreover, we detected and quantified Fetuin B signals using immunofluorescence staining and immunoblotting in mouse embryos treated with MG132. We found that MG132 treatment leads to Fetuin B accumulation in mouse embryos (Figure 5C, Figures S8A-B), supporting that Fetuin B is degraded by proteasomes.

**Figure 5.**
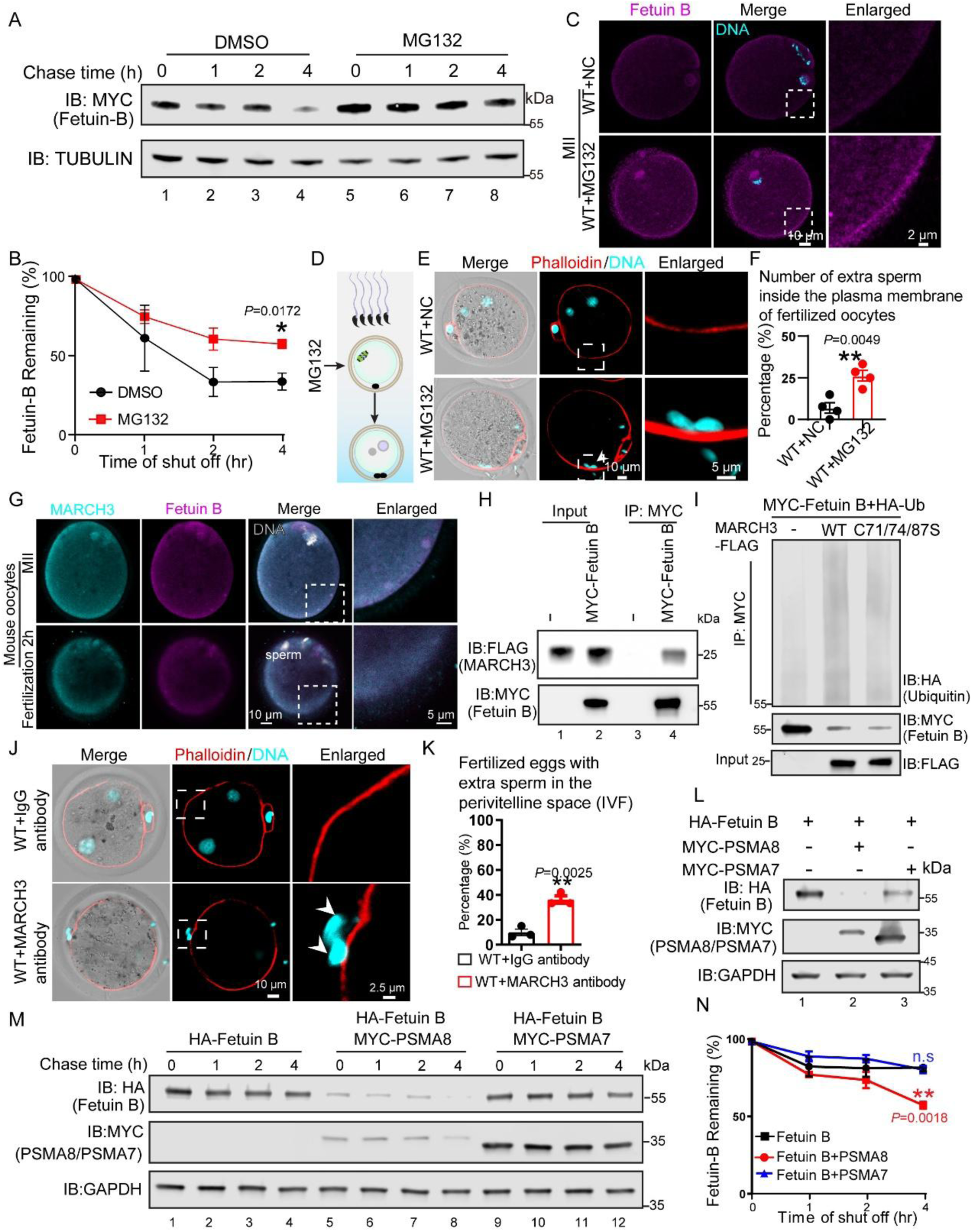
MARCH3 mediates polyubiquitination and degradation of Fetuin. **B** (A) MG132 treatment leads to the accumulation of Fetuin B. A cycloheximide chase (CHX) assay was performed to examine Fetuin B in HEK293T cells treated with DMSO or MG132. Samples were collected at 0, 1, 2, and 4 hours after CHX addition. Fetuin B protein levels were detected by immunoblotting with a MYC antibody. GAPDH served as a loading control. (B) Quantification of Fetuin B levels in (A). Protein levels of Fetuin B were quantified using Odyssey software. (n = 3 independent experiments). Data are presented as the mean ± SEM. Two-tailed unpaired Student’s t test, *p < 0.05. (C) Representative images showing the localization of Fetuin B in control and MG132-treated mouse oocytes. (D) Schematic of sample collection in DMSO (negative control, NC) and MG132 treated embryos. (E) Representative images of mouse embryos stained with Phalloidin (red) and DAPI (cyan) in DMSO and MG132 treated embryos after 6 hours post IVF. Arrows indicate extra sperm inside the plasma membrane of fertilized oocytes. (F) Percentage of fertilized oocytes with extra sperm inside the plasma membrane was quantified in DMSO and MG132 treated embryos. WT+NC, 6.98% ± 3.12% (n = 4 independent experiments, total oocytes = 89); WT+MG132, 26.28% ± 3.18% (n = 4 independent experiments, total oocytes = 64). Data are presented as the mean ± SEM. Two-tailed unpaired Student’s t test, **p < 0.01. (G) Representative images showing the localization of Fetuin B and MARCH3 in mouse oocytes and embryos at the indicated developmental stages. (H) Fetuin B physically interacts with MARCH3. MYC-Fetuin B and MARCH3-FLAG (or empty vector) were co-transfected into HEK293T cells. 24 hours after transfection, cells were collected for immunoprecipitation with an anti-MYC antibody, followed by analysis using anti-FLAG and anti-MYC antibodies. (I) MARCH3 but not its E3 ligase-inactive mutants promotes the polyubiquitination of Fetuin B. HEK293T cells were transfected with MYC-Fetuin B, MARCH3-FLAG, MARCH3 C71/74/87S-FLAG, or HA-Ub. 24 hours after transfection, cells were collected for immunoprecipitation with an anti-MYC antibody, followed by analysis with anti-HA, anti-FLAG and anti-MYC antibodies. (J) Representative images of mouse embryos stained with Phalloidin (red) and DAPI (cyan), Embryos are from mice receiving control IgG antibody or MARCH3 antibody 6 hours post IVF. Arrows indicate extra sperm in the perivitelline space. (K) Percentage of fertilized oocytes with extra sperm in the perivitelline space was quantified in (J). WT+ IgG antibody, 9.87% ± 2.60% (n = 3 independent experiments, total oocytes = 75); WT+ MARCH3 antibody, 36.23% ± 2.93% (n = 3 independent experiments, total oocytes = 52). Data are presented as the mean ± SEM. Two-tailed unpaired Student’s t test, **p < 0.01. (L) Overexpression of PSMA8 promotes the degradation of Fetuin B more effectively than PSMA7. (M) PSMA8 accelerates Fetuin B degradation. A CHX assay of Fetuin B was performed in MARCH3 and PSMA8 or PSMA7 expressed HEK293T cells. Samples were taken at 0, 1, 2, and 4 h after the addition of CHX. The Fetuin B, PSMA8 and PSMA7 protein levels were detected by immunoblotting using HA and MYC antibody. GAPDH served as a loading control. (N) Quantification of the Fetuin B levels in (M). The protein level of Fetuin B was quantified using Odyssey software. (n = 3 independent experiments). Data are presented as the mean ± SEM. Two-tailed unpaired Student’s t test, **p < 0.01, n.s., non-significant.

We next investigated whether proteasome dysfunction increases embryo susceptibility to polyspermy. Extra sperm were found inside the plasma membrane of MG132-treated oocytes during fertilization (Figures 5D-E, Figure S8C). The percentage of fertilized oocytes with polyspermy (≥3PN) significantly increased in those receiving MG132 compared to the control group (Figure 5F), indicating disruption of proteasome function results in polyspermy.

To determine whether the proteasome’s role in preventing polyspermy is conserved between humans and mice, we tested proteasome inhibitors on human oocyte fertilization. Human MII oocytes were treated with MG132 for 1 hour, and then combined with human sperm for 8 hours. Immunofluorescence staining showed that extra sperm were found inside the plasma membrane of MG132-treated human oocytes (Figures S8D-E), suggesting functional conservation of the proteasome.

It was recently reported that zygotes deficient in the ubiquitin E3 ligase MARC-3 (the homolog of murine MARCH3) show a polyspermy phenotype in *Caenorhabditis elegans*.^41^ To investigate potential functional relationship(s) between MARCH3 and Fetuin B during fertilization, we examined the subcellular localization of these two proteins in oocytes and embryos. MARCH3 and Fetuin B are both localized in the cytoplasm of oocytes and embryos in both humans and mice (Figure 5G, Figure S8F). Furthermore, we constructed MYC-Fetuin B and FLAG-MARCH3 plasmids, which we transfected into HEK293T cells. Co-immunoprecipitation (Co-IP) analysis revealed that MARCH3 physically interacts with Fetuin B (Figure 5H, Figure S8G). In addition, overexpression of MARCH3 in HEK293T cells mediated the polyubiquitination of Fetuin B (Figure S8H). Multiple E3 ligase-inactive mutants of MARCH3 have been reported, including C71S, C74S, and C87S.^42^ We generated a triple mutant MARCH3 variant, which we designated as MARCH3 C71/74/87S. We found that MARCH3, but not its E3 ligase-inactive mutants, mediates the polyubiquitination of Fetuin B in HEK293T cells (Figure 5I), confirming that MARCH3 mediates the polyubiquitination of Fetuin B.

To further elucidate the functional role of MARCH3 in fertilization, we performed IVF with WT oocytes in the presence of MARCH3 antibody or IgG antibody during fertilization. We found that early developmental arrest occurred with MARCH3 antibody but not control IgG antibody. (Figures S8I-J). Additionally, extra sperm were observed in the perivitelline space of MARCH3-blocked pronuclei and two-cell embryos; no extra sperm were observed in those receiving control IgG antibody (Figure 5J, Figure S8K). Quantitative analysis revealed a significantly higher percentage of embryos with extra sperm in the perivitelline space in the presence of MARCH3 antibody compared to the control group (Figure 5K). These results indicate that blocking MARCH3 leads to zona pellucida block failure, suggesting that MARCH3 prevents extra sperm penetrating zona pellucida by mediating the polyubiquitination and degradation of Fetuin B during fertilization.

### The newly assembled chimeric proteasome promotes the degradation of Fetuin B

Recalling our observation that both 20S CP components and 19S RP components are present in oocytes, with the exception of PSMA8, why are sperm 20S CPs necessary to assemble with oocyte 19S RPs for proteasome assembly during fertilization? We propose that sperm 20S CPs integrated with oocyte 19S RPs to form an active chimeric proteasome, which facilitates the timely degradation of maternal proteins (e.g., Fetuin B) to prevent polyspermy during fertilization.

To further elucidate the function of the newly assembled chimeric proteasome during fertilization, we initially characterized Fetuin B levels in HEK293T cells that expressed either the testis-specific proteasome subunit PSMA8 or PSMA7. We found PSMA8 expression significantly decreased Fetuin B levels compared with PSMA7 or no expression (Figure 5L). In addition, chase assays indicate that the decrease in Fetuin B levels is due to increased proteasomal degradation rather than reduced synthesis (Figures 5M-N). Collectively, these results demonstrate that PSMA8 expression promotes the assembly of testis-specific proteasome, which exhibits higher Fetuin B degradation activity compared to conventional proteasome.

## Discussion

Numerous maternal factors have been reported to prevent polyspermy and ensure successful embryonic development.^5,10,17^ Traditionally, sperm cells have been regarded merely as vehicles for delivering the paternal haploid genome to the oocyte. Consequently, the paternal factors that trigger polyspermy have long been overlooked. In this study, our findings provide genetic, cellular, and biochemical evidence that reveals 20S CPs stored in the sperm head are integrated with oocyte-derived 19S RPs to provide newly assembled chimeric proteasomes during fertilization. The proteasomes are then distributed to the oocyte and, with the help of MARCH3, promote Fetuin B degradation. This degradation leads to the cleavage of ZP2 and the hardening of the zona pellucida, ultimately preventing polyspermy in embryos (Figure 6). We have uncovered a molecular mechanism involved in mammalian fertilization that enhances our understanding of paternal factors that prevent polyspermy. Previous investigations have revealed that paternal factors (e.g., sperm DNA, RNA, proteins, and centrioles) contribute to fertilization and early embryonic development, though their functional scope in post-fertilization events remains partially characterized.^23,24^ Notably, Castillo *et al*. (2018) highlighted that a subset of sperm proteins is crucial “for fertilization and beyond,” suggesting broader roles than previously recognized. Our study extends this paradigm by demonstrating that paternal factors stored in sperm nuclear vacuoles may further influence embryogenesis and offspring development.

**Figure 6.**
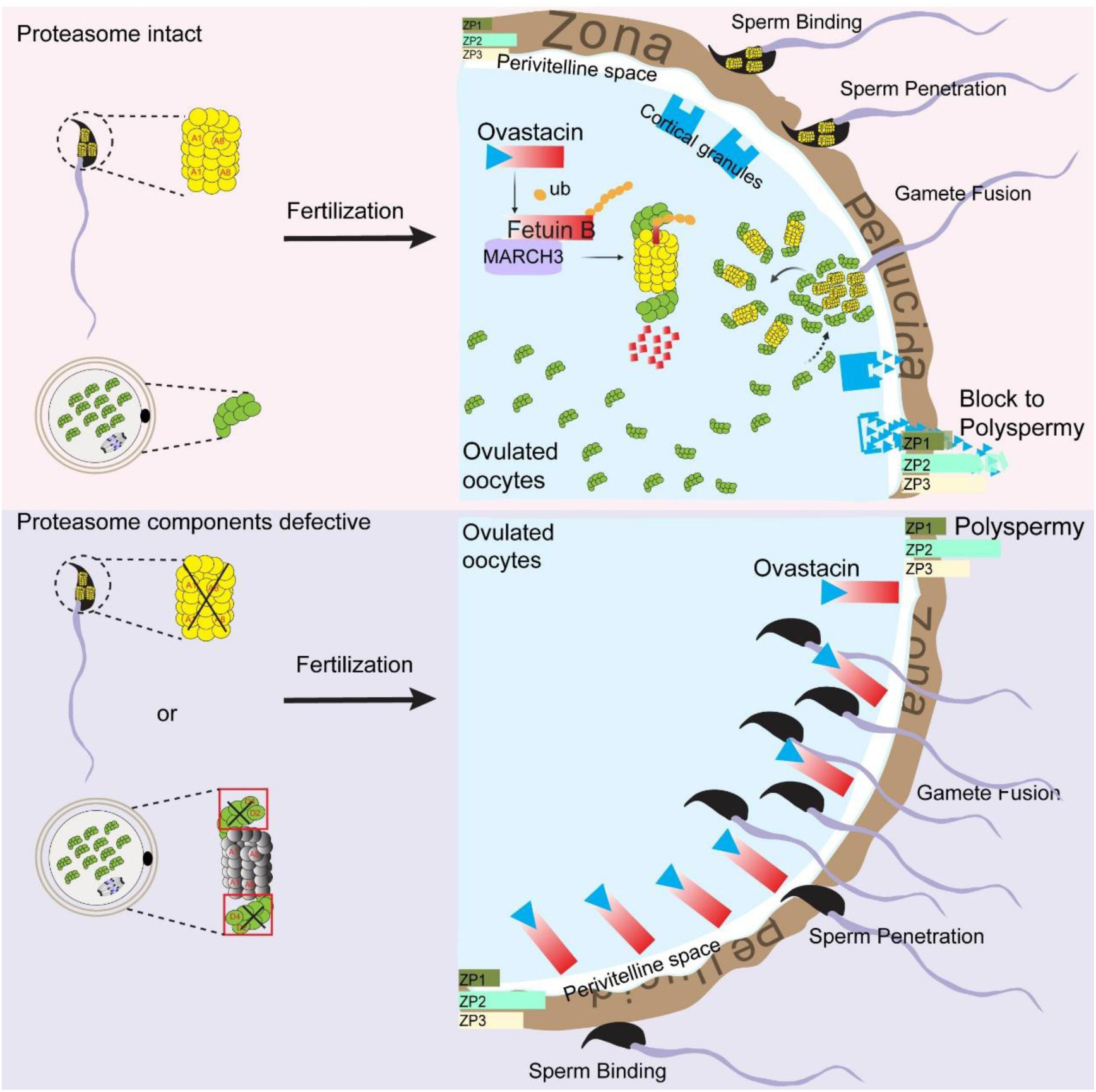
Separately prestored proteasome components prevent polyspermy during fertilization. Model of proteasome components function in preventing polyspermy during mammalian fertilization. In normal sperm and oocytes (top), Prestored 20S CPs in the sperm head were integrated with oocyte-derived 19S RP components to generate newly assembled chimeric proteasome during fertilization. The proteasomes are then distributed to the oocyte and, with the help of MARCH3, promotes Fetuin B degradation. This degradation leads to the cleavage of ZP2 and zona pellucida hardening, ultimately preventing polyspermy in embryos. In sperm with defective 20S CP components or oocytes with defective 19S RP components (bottom), new chimeric proteasomes do not form, leading to the accumulation of Fetuin B. The accumulation of Fetuin B inhibits Ovastacin, resulting in uncleaved ZP2, which prevents the hardening of the zona pellucida, ultimately causing polyspermy in embryos.

Polyspermy prevention in mammals occurs very rapidly, and the membrane block is established within 30-45 minutes after fertilization.^5^ At this stage, zygotic genes have not been activated, so transcription has not yet occurred. Even with prestored maternal mRNA in the oocyte, the rate of translation and protein modification cannot meet the requirements necessary for a rapid response to block polyspermy. Thus, prestored proteins delivered from the sperm to the oocyte might facilitate this rapid response. Since this process must occur after fertilization, factors prestored in sperm nuclear vacuoles might trigger this process. Upon fertilization, oocyte-derived 19S RPs integrate with testis-specific 20S CPs to assemble chimeric proteasomes, which are then quickly distributed within oocytes to trigger a signaling cascade that eventually blocks polyspermy (Figure 3D). Thus, prestored 20S CPs in sperm and the assembly of chimeric proteasomes after fertilization, which initiate precisely timed protein degradation essential for fertilization.

Recent studies have shown that 20S CPs and 19S RPs do not always function in concert.^43–45^ In neurons, 20S CPs and 19S RPs have been found to be differentially distributed in neuronal subcellular compartments, with an abundance of free 19S RPs near synapses to regulate neuronal activity.^43^ Different from previous discoveries, our findings suggest prestored proteasome components in oocytes and in sperm might represent a strategy to prevent the premature degradation of Fetuin B, thereby preventing inappropriate hardening of the zona pellucida in MII oocytes before fertilization.

Proteasome complexes exist in various configurations in eukaryotes, and even tissue-specific proteasome components have emerged.^46^ Besides the most common 19S RPs, two additional proteasome regulatory particles have been identified that can activate 20S CPs. The 11S proteasome regulatory particle (PA28α, PA28β, and PA28γ) functions to degrade oxidized proteins, process antigens, and strengthen immune responses against malignant cells.^47,48^ The PA200 regulatory particle functions in the acetylation-dependent degradation of histones during DNA repair and histone-protamine exchange.^34^ In immune cells, the 20S immunoproteasome preferentially incorporates catalytic subunits β1i/LMP2, β2i/MECL-1, and β5i/LMP7, replacing corresponding constitutive subunits.^49^. The testis-specific proteasome contains α4s/PSMA8 rather than α4/PSMA7, which has been reported to be essential for proper meiotic exit in spermatocytes.^31–33^ Our work shows, for the first time, that such a replacement boosts proteasome activity to degrade Fetuin B in zygotes and suggests that paternal proteasome components are essential to degrade maternal proteins and block polyspermy.

The molecular composition of 20S CPs is conserved across eukaryotes, with subtle subunit replacements in tissue-specific homologs.^50^ In mammalian testis, PSMA8 is an α-type subunit with high homology to α4 (PSMA7),^31^ and the origin of this testis-specific subunit can be traced back to invertebrates (Figure S9), suggesting that its function may be evolutionary conserved across invertebrates to mammals. The function of MARCH3 in preventing extra sperm penetrating zona pellucida is also conserved from worm to mammals (Figure S10). Thus, although we have demonstrated that prestored 20S CPs in sperm heads are involved in preventing polyspermy in human and mouse, this mechanism might also apply to a broad range of eukaryotic organisms.

## METHODS

### Reagent or resource

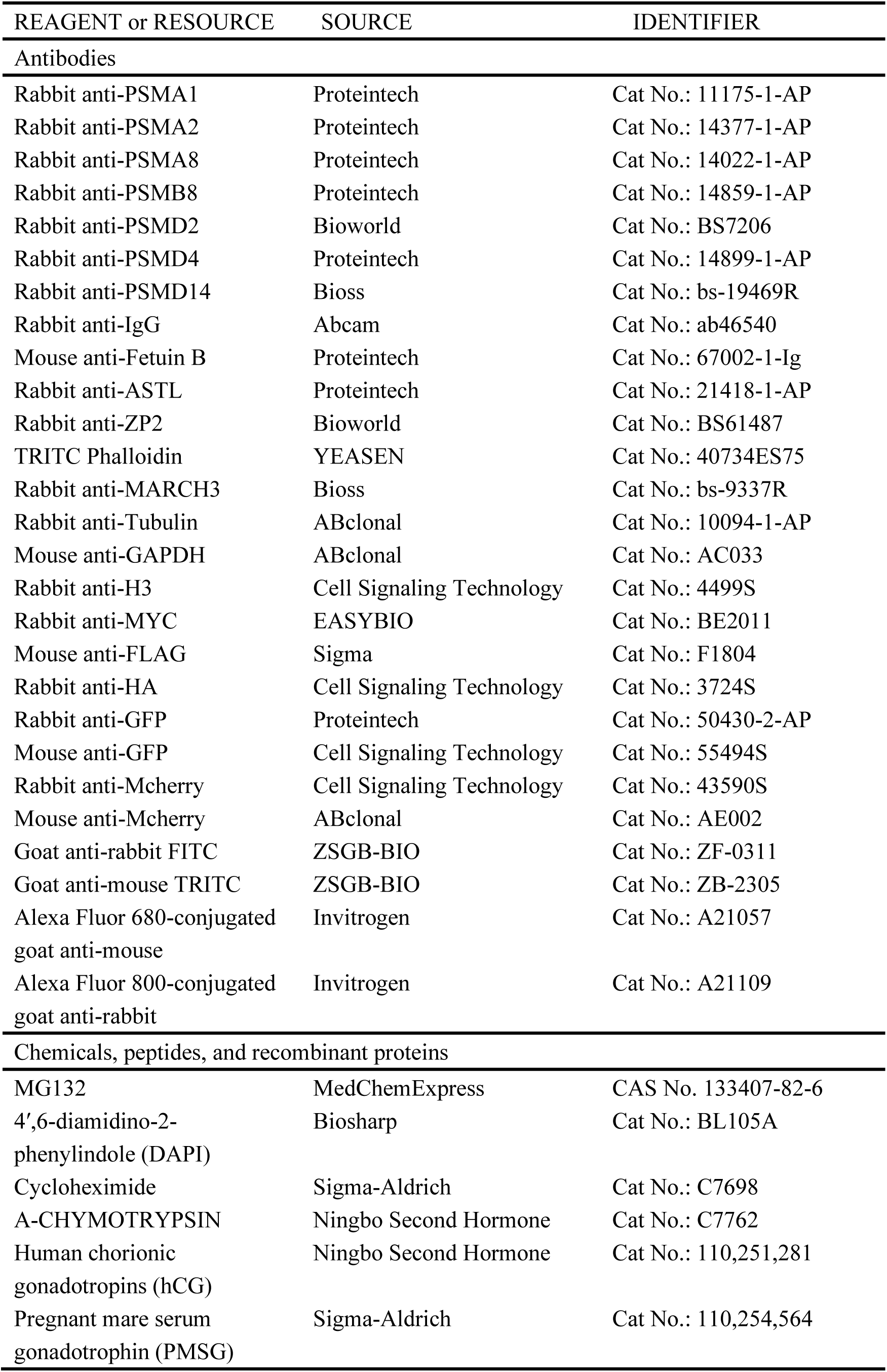
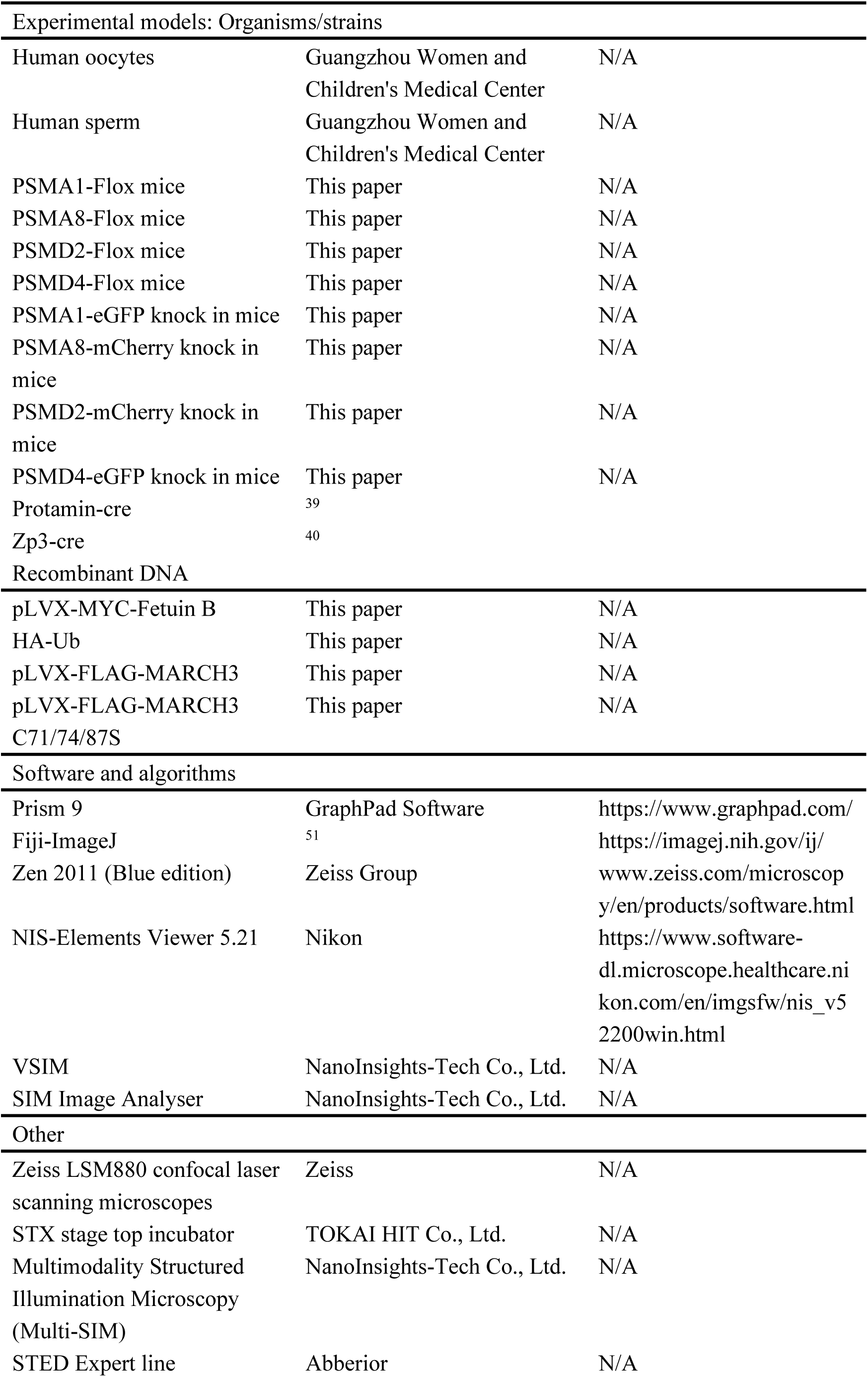
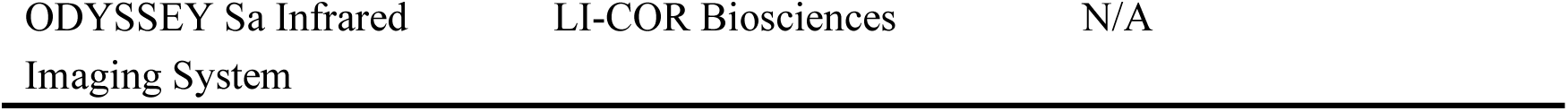

### Animals

*Psma1^Flox/Flox^*, *Psma8^Flox/Flox^*, *Psmd2^Flox/Flox^* or *Psmd4^Flox/Flox^* mice were generated using the CRISPR/Cas9 system. Briefly, LoxP sites were inserted into the *Psma1* genomic region, which includes exons 5, 6, and 7, using recombineering to generate *Psma1^Flox/+^* mice. Similarly, LoxP sites were inserted into the *Psma8* genomic region, which includes exon 2, to generate *Psma8^Flox/+^* mice; into the *Psmd2* genomic region, which includes exons 4-8, to generate *Psmd2 ^Flox/+^*mice; and into the *Psmd4* genomic region, which includes exons 2 and 3, to generate *Psmd4 ^Flox/+^* mice. These mice were maintained on a B6/129 genetic background. Mice lacking *Psma1*, *Psma8*, *Psmd2,* or *Psmd4* in their oocytes were generated by crossing *Psma1^Flox/Flox^*, *Psma8^Flox/Flox^*, *Psmd2^Flox/Flox^* , or *Psmd4^Flox/Flox^* mice with *Zp3* promoter-mediated Cre transgenic mice^40^. Similarly, mice lacking *Psma1*, *Psma8*, *Psmd2,* or *Psmd4* in their spermatozoa were generated by crossing these mice with *Protamin1* promoter-mediated Cre transgenic mice^39^. The *Prm-Cre* knock-in mouse line was generously provided by Dr. Wei Li from the State Key Laboratory of Stem Cell and Reproductive Biology, Institute of Zoology, Stem Cell and Regenerative Medicine Innovation Institute, Chinese Academy of Sciences.

To generate knock-in mice, the donor vector containing the “3xEAAAK-EGFP” cassette, the gRNA targeting mouse *Psma1* or *Psmd4* gene, and Cas9 mRNA were co-injected into fertilized mouse eggs to produce targeted knock-in offspring. The donor vector containing the “3xEAAAK-mCherry” cassette, the gRNA targeting mouse *Psma8* or *Psmd2* gene, and Cas9 mRNA were similarly co-injected to generate targeted knock-in offspring. F0 founder animals were identified by PCR followed by sequence analysis and were bred to wild-type mice to test germline transmission and produce F1 animals. The mice were maintained under specific pathogen-free (SPF) conditions in a controlled environment with temperatures of 20-22°C, 12/12-hour light/dark cycles, 50-70% humidity, and food and water provided *ad libitum*. Ethical approval for all animal experimental procedures was provided by the institutional animal care and use committee of Guangzhou Medical University (GZ2024-116).

### Superovulation and fertilization

For superovulation, female mice were injected intraperitoneally with 5 IU of pregnant mare serum gonadotrophin (PMSG, 110,254,564, Ningbo Second Hormone Company). This was followed 46-48 hours later by an injection of 5 IU human chorionic gonadotropin (hCG, 110,251,281, Ningbo Second Hormone Company). After an additional 13 hours, oocyte/cumulus masses were surgically removed from the ampulla of the oviducts. The oocytes were then harvested and used for immunofluorescent assay after digestion with 0.3% hyaluronidase (Sigma-Aldrich) to remove cumulus cells. To obtain fertilized eggs (zygotes), 8-week-old female mice were mated with 10-week-old WT males. Successful mating was confirmed by the presence of vaginal plugs. E0.5 (zygotes) and E1.5 (2-cell) embryos were flushed out of the oviducts and used for immunofluorescent assay.

### *In vitro* fertilization and Sperm binding assay

Sperm was released from the cauda epididymides of 10-week-old male mice into 200 μL of human tubal fluid medium (HTF, Sudgen Biotechnology, China) and capacitated for 1 hour at 37°C in a 5% CO_2_ incubator. Meanwhile, ovulated oocytes were harvested as previously described. These oocytes were then cultured in 100 μL of HTF, covered with mineral oil (M5310, Sigma-Aldrich), and left to mature at 37°C in a 5% CO_2_ atmosphere for *in vitro* fertilization. For fertilization, 4 × 10^5/mL capacitated sperm were incubated with the ovulated oocytes in HTF for 3 hours. Fertilized oocytes were then washed in fresh HTF drops to separate them from abnormal oocytes, and the process took place at 37°C on a TPiE-SMZR (TOKAI HIT) constant temperature plate. Successful fertilization was confirmed by the presence of pronuclei after 6 hours.

For the sperm binding assay, 4 × 10^5/mL capacitated sperm were incubated with ovulated oocytes in 100 μL HTF for 1 hour at 37 °C in a humidified atmosphere of 5% CO_2_. Sperm binding to ovulated oocytes and 2-cell embryos were observed using capacitated sperm. Negative controls were generated by washing with a wide-bore pipette to remove all but two to six sperm on normal two-cell embryos. Samples were fixed in 4% PFA and 0.5% Triton X-100 for 20 minutes and then stained with 4’,6-diamidino-2-phenylindole (DAPI) to quantify sperm number. Bound sperm were quantified from z-projections acquired by a confocal microscope.

### Human oocyte collection and manipulation

Human oocytes and sperm were obtained from Guangzhou Women and Children’s Medical Centre, Guangzhou, China. Human oocytes were obtained from donors undergoing intracytoplasmic sperm injection treatment for male infertility. Unmatured oocytes were selected from donated cells cultured with *in vivo* maturation media (ART-1600) and allowed to mature at 37°C in a 6% CO_2_ atmosphere, and then randomly assigned to control and experimental groups for *in vitro* fertilization. For fertilization, 1×10^5/mL human sperm were incubated with the matured oocytes in HTF medium (Vitrolife, 10136) for 6 hours, followed by collection for immunofluorescence staining. This study was approved by the Medical Ethics Committee of Guangzhou Women and Children’s Medical Center (ID: 2024126A01). The procedures were conducted in accordance with the International Ethical Guidelines for Research Involving Human Subjects, as outlined in the Helsinki Declaration. The procedures were conducted in accordance with the International Ethical Guidelines for Research Involving Human Subjects, as outlined in the Helsinki Declaration.

### Zona Pellucida Digestion

Zona pellucida digestion was performed as previously reported using a 0.2 mg/mL α-chymotrypsin (10 IU/mL) solution.^52^ Briefly, the percentage of zona-free oocytes was plotted against time. T50 represents the time point at which 50% of oocytes became zona-free.

### Histological analysis

Mouse testes were fixed in 4% paraformaldehyde (PFA, P1110, Solarbio, China) overnight at 4°C, dehydrated in an ethanol series, and embedded in paraffin. Next, 5-μm sections were cut with a microtome. Following deparaffinization and rehydration, the sections were stained with hematoxylin and eosin (G1100, Solarbio, China) to observe the spermatogenesis process. Images were collected using a Nikon inverted microscope with a charge coupled device (CCD) (Nikon, Eclipse Ti-S, Tokyo, Japan).

### Immunofluorescence

For oocyte and embryo immunostaining, oocytes or embryos were fixed and permeabilized in 4% PFA and 0.5% Triton X-100 for 20 minutes at room temperature. After blocking with 1% bovine serum albumin (BSA, AP0027, Amresco, USA) in PBS, oocytes were incubated with primary antibodies diluted in 1% BSA overnight at 4°C. Following three washes with PBS, oocytes were labeled with secondary antibodies for 1 hour at 37°C and then stained with DAPI for 5 minutes. Finally, oocytes were mounted on glass slides and observed with a Leica SP8 microscope (Wetzlar, Germany) or a Nikon AXR microscope (Tokyo, Japan).

For spermatozoa immunostaining, spermatozoa were released from the cauda epididymis of WT or proteasome-knockout male mice and incubated in PBS at 37°C for 30 minutes. They were then spread on glass slides for morphological observation or immunostaining. Following air drying, the spermatozoa were fixed in 4% PFA for 10 minutes and subsequently permeabilized in 0.5% Triton X-100 for 20 minutes at room temperature. The slides were washed three times with PBS and then blocked with 5% BSA for 30 minutes at room temperature. Primary antibodies were added to sections, incubated at 4°C overnight, and then followed by incubation with secondary antibody. Finally, the nuclei were stained with DAPI, and images were captured using a confocal microscope.

### Super-resolution imaging

Super-resolved images were acquired using a Multimodality Structured Illumination Microscopy (Multi-SIM) or a Stimulated emission depletion (STED) Expert line system. Embryos incubated with primary antibody were washed three times in 0.5% Triton X-100/PBS (PBT) before incubation with secondary antibodies (Abberior STAR RED and Abberior STAR ORANGE). STED images were acquired using a STEDYcon mounted on an inverted Nikon microscope, which was equipped with pulsed diode lasers at 640 nm and 561 nm, a 775 nm laser, a QUAD Scanner (80 µm×80 µm, 100×/1.4 NA oil), and three photon-counting avalanche photodiodes (APDs).

Multi-SIM images of sperm were acquired using a Multi-SIM imaging system (NanoInsights) equipped with a 100×/1.49 NA oil objective (Nikon CFI SR HP Apo), solid-state lasers (561 nm, 640 nm), and a sCMOS (Complementary Metal-Oxide-Semiconductor) camera (Photometrics Kinetix). Serial Z-stack sectioning was performed for the Stacked Slices-SIM mode. SIM image stacks were reconstructed using SI-Recon 2.23.3 (NanoInsights) with the following settings: pixel size of 30.6 nm, channel-specific optical transfer functions, a Wiener filter constant of 0.01 for 2D mode, and the discarding of negative intensity backgrounds. Reconstructed SIM images were then denoised using a total variation (TV) constraint.

### Fertility testing

For female fertility testing, 5 individually *Psma1^flox/flox^* and *Zp3-Psma1^-/-^*, *Psma8^flox/flox^* and *Zp3-Psma8^-/-^*, *Psmd2^flox/flox^* and *Zp3-Psmd2^-/-^*, or *Psmd4^flox/flox^* and *Zp3-Psmd4^-/-^* female mice at the age of 8 weeks were continuously mated with 10 weeks old fertile males. For male fertility testing, 3 individually *Psma1^flox/flox^* and *Prm-Psma1^-/-^*, *Psma8^flox/flox^* and *Prm-Psma8^-/-^*, *Psmd2^flox/flox^*and *Prm-Psmd2^-/-^*, or *Psmd4^flox/flox^* and *Prm-Psmd4^-/-^* male mice at the age of 8 weeks were caged with two wild-type female mice, and their vaginal plugs were checked every morning. The plugged females were separated and caged individually, and pregnancy outcomes were recorded. Females that did not generate offspring at 22 days post conception were scored as not being pregnant. The numbers of pups and average litters were recorded for the plugged females.

### Time-lapse live imaging experiments

Capacitated sperm were incubated with ovulated oocytes in HTF and then transferred into a well with two spacer layers within a 4-well 0.05 mm chamber (SUNJin Lab #IS204) affixed to a clean glass slide. A glass coverslip was carefully placed on top. The live fertilized oocytes were imaged using a Nikon AXR confocal imaging system equipped with an STX stage top incubator (TOKAI HIT). The imaging process took place in a controlled environment with a 5% CO2 atmosphere at 37°C.

### Proximity ligation assay

Protein-protein interactions were detected using the DuoLink® In Situ Red Starter Kit (Mouse/Rabbit) (DUO92101, Sigma-Aldrich). PSMA1-eGFP capacitated sperm were co-incubated with ovulated oocytes in HTF medium for 1.5 hours. Following fertilization, the fertilized oocytes were washed in fresh HTF droplets to remove abnormal oocytes. The embryos were then fixed and permeabilized in 4% paraformaldehyde (PFA) and 0.5% Triton X-100 for 20 minutes at room temperature. Subsequently, the embryos were blocked with Duolink® Blocking Solution for 1 hour at 37°C. Primary antibodies against GFP and PSMD2 were applied to detect PSMA1 and PSMD2, followed by overnight incubation at 4°C. The samples were then washed with 1× Wash Buffer A and incubated with two PLA probes (diluted 1:5 in antibody diluent) for 1 hour. Next, the Ligation-Ligase solution was applied for 30 minutes, followed by the Amplification-Polymerase solution for 100 minutes in a pre-heated humidified chamber at 37°C. Prior to imaging, the embryos were washed with 1× Wash Buffer B and mounted on coverslips using Duolink® In Situ Mounting Medium containing DAPI. Fluorescence images were acquired using a Nikon AXR microscope (Tokyo, Japan).

### Western blot analysis

A total of 200 mouse oocytes were collected and transferred to 10 μl of 2% SDS sample buffer, then boiled for 10 minutes at 95°C for subsequent immunoblotting. To prepare cell protein extracts, cells were suspended in cold RIPA buffer (R0010, Solarbio) supplemented with a protein inhibitor cocktail (Roche Diagnostics, 04693116001, Rotkreuz, Switzerland) and 1 mM phenylmethyl sulfonyl fluoride (PMSF, 0754, Amresco). After homogenization and transient sonication, cell extracts were incubated on ice for 30 minutes. The samples were then centrifuged at 12,000 × rpm for 15 minutes at 4°C, and the supernatant was transferred to a new tube for immunoblotting. Protein lysates were separated via SDS-PAGE and electro-transferred to a nitrocellulose membrane. The membranes were blocked in PBS containing 5% skimmed milk for 1 hour at room temperature, followed by overnight incubation at 4°C with primary antibody. Afterward, membranes were incubated with secondary antibody for 1 hour at room temperature. Finally, membranes were washed three times in PBS buffer and scanned using an ODYSSEY Sa Infrared Imaging System (LI-COR Biosciences, Lincoln, NE, USA).

### Cell culture and plasmid transfection

HEK293T cells were cultured in DMEM (Invitrogen) supplemented with 10% fetal bovine serum (Hyclone) and 1% penicillin-streptomycin solution (Gibco) at 37°C in a 5% CO2 incubator. cDNAs encoding mouse Fetuin B and MARCH3 were subcloned into pLVX vectors with MYC or FLAG tags for subsequent immunoprecipitation. MARCH3 C71/74/87S mutant was created using the KOD-Plus-Neo (KOD-401) enzyme. HEK293T cells were transfected with MYC-Fetuin B, HA-Ub, and either FLAG-MARCH3 or the MARCH3 mutant plasmids using LipoMax DNA Transfection Reagent (Sudgen Biotechnology, China) according to the manufacturer’s instructions for subsequent ubiquitination assays.

### Co-Immunoprecipitation

At 48 hours post-transfection, cells were lysed in a lysis buffer containing 50 mM Tris-HCl (pH 7.5), 150 mM NaCl, 10% glycerol, and 0.5% NP-40, with protease inhibitors added immediately before use. After sonication and high-speed centrifugation of the cell lysates, the supernatant was incubated with primary antibody overnight at 4°C. Subsequently, samples were incubated with protein A-Sepharose (GE, 17-1279-03) for 2 hours at 4°C. Following this incubation, precipitates were washed three times with IP buffer (20 mM Tris, pH 7.4, 2 mM EGTA, and 1% NP-40), and bound proteins were analyzed with immunoblotting.

### Cryo ET sample preparation

Quantifoil R2/1 EM grids were pre-coated with a 20 nm carbon film using EM ACE200 (Leica), followed by glow discharging for 45 s in a GATAN Model 950 Advanced Plasma System. Human sperm samples were prepared by collecting the lower milky layer after standing for 40 minutes. Mouse sperm samples were diluted to 6-8×10^7^ cells/mL in PBS buffer. Before preparation, glycerol was added to a final concentration of 10% as a cryoprotectant. Aliquots of 4 μL were pipetted onto the treated EM grids, blotted 3∼4 times from the reverse side for 8 seconds each, and then plunge-frozen into a liquid nitrogen-cooled ethane-propane mixture using a manual plunger.

### Lamellae preparation

Lamellae were prepared using an Aquilos2 Cryo-FIB system (Thermofisher Scientific) as described.^53^ In brief, the EM grid was treated with 30 s of gas injection for platinum deposition, followed by 15 s of platinum sputtering at 30 mA to create protective layers. The stage was then tilted to 20° for milling. Rough milling was performed with a 30 kV Ga^2+^ ion beam at 1 nA, with the current gradually reduced (0.3 nA, 0.1 nA, 50 pA, 30 pA) near the target region to avoid radiation damage. Final polishing thinned the lamellae to less than 200 nm at 10 pA.

### Cryo-ET data acquisition and reconstruction

Lamellae were loaded into a 300kV Titan Krios TEM (Thermo Fisher Scientific) equipped with a post-GIF K3 direct detector (Gatan). Low-magnification TEM images (2600×) of lamellae were acquired with 20 s exposure time. Tilt series were acquired using SerialEM software,^54^ employing a dose-symmetric tilt scheme covering angles from -40° to +60° (relative to the pre-tilt angle) in 2° increments.^55^ Images were captured at an object pixel size of 2.64 Å and a defocus range of - 4 to - 6 μm in counting mode. The total electron dose was 120 e^−^/Å², with each tilt receiving approximately 2.35 e^−^/Å², distributed over a movie of 10 frames. In total, 60 tomograms were obtained from 20 lamellae.

The individual projection images were motion-corrected by MotionCor2 software.^56^ Tilt series were aligned using the patching tracking method in IMOD 4.11.3 software,^57^ CTF estimation and back projection reconstruction were established in Warp.^58^

### Template matching and subtomogram averaging

Initially, particles were manually selected from a single tomogram. Subsequent alignment and averaging revealed a 20S CP structure. This map served as a reference for template matching across all tomograms. Template matching was conducted on downsampled tomograms (binning factor of 6) using pyTOM.^59^ After manual validation, a total of 6,698 particles were identified from 15 tomograms.

For subtomogram averaging, particle coordinates were input into Warp to extract full-size subtomograms.^60^ Subsequent classification and refinement were performed using RELION/3.1.1.^61^ Firstly, a global 3D classification without masking was performed, resulting in the exclusion of 12% of poorly aligned particles. The remaining particles were refined to reach accurate Euler angles. These well-aligned particles were then subjected to further classification without additional alignment (with C7 symmetry and extended mask applied). Ultimately, 2,906 particles were refined with C7 symmetry, resulting in a map with a resolution of 15.6 Å (Figure S1B). Subsequently, the 20S CP were placed back into the tomogram according to the refined coordinates and Euler angles, and then visualized in ChimeraX.^62^

### Chase assays

HEK293T cells were plated one day before the experiment, and cycloheximide (R750107; CHX; Sigma-Aldrich) was added to the culture at a concentration of 100 μg/ml to block new protein synthesis. Samples were taken at 0, 1, 2, and 4 hours after the addition of CHX for 2 hours. Protein levels were determined by immunoblotting. To distinguish proteasomal degradation, the cells were treated with 20 μM MG132 (sc-201270; Santa Cruz) for 12 hours to inhibit proteasomal degradation.

### Phylogenetic tree-building

To construct a phylogenetic tree, multi-sequence alignment of the different genes was first performed using Mafft v7.505. Subsequently, the trimAl v1.5.rev0 software was employed to remove ambiguous regions and gaps from the alignment, retaining only highly conserved regions. Next, FastTree v2.1.11 was used to generate the phylogenetic tree using the maximum likelihood method. The default JTT+T model was selected as the protein substitution model, and the bootstrap resampling value was set to 1000.

### Statistical analysis

Statistical analyses were conducted using GraphPad PRISM version 9. All experiments were repeated at least three times and all data were expressed as mean ± SEM or mean ± SD. The statistical significance of the differences between the mean values for the different genotypes was measured by two-tailed unpaired student’s t test. The data were considered statistically significant when the P value was less than 0.05 (*), 0.01 (**), 0.001 (***), or 0.0001 (****).

## ACKNOWLEDGMENTS

This work was supported by the National Key R&D Program of China (2024YFC2706802), the National Natural Science Foundation of China (32400709, 32230029, 32270898, 32400714) , the Postdoctoral Fellowship Program of CPSF under Grant Number GZC20251824, the China Postdoctoral Science Foundation (2024M760638, 2024M760640).

## AUTHOR CONTRIBUTIONS

W.L., Q.G., R.J. and X.G. conceptualized the study and designed the experiments and data analysis methods. L.W. and C.L. conducted all experiments and performed the data analysis.

X.W. established the cryo-ET acquisition and analysis workflow. L.W., Q.Z., Y.C., H.W. and X.H. developed conditions for mouse oocyte vitrification and expansion microscopy. Y.G., R.J., W.L. and C. S. provided input on experimental design and analysis. S.L. and L.S. oversaw the collection of human oocytes. L.W. and W.L. drafted the manuscript and prepared the figures.

## INCLUSION AND DIVERSITY

We support inclusive, diverse, and equitable conduct of research.

