## Supplementary Figure1-10 for "Separately prestored proteasome components to prevent polyspermy"

**Supplementary Materials:**

**Supplementary Figures and Figure Legends**

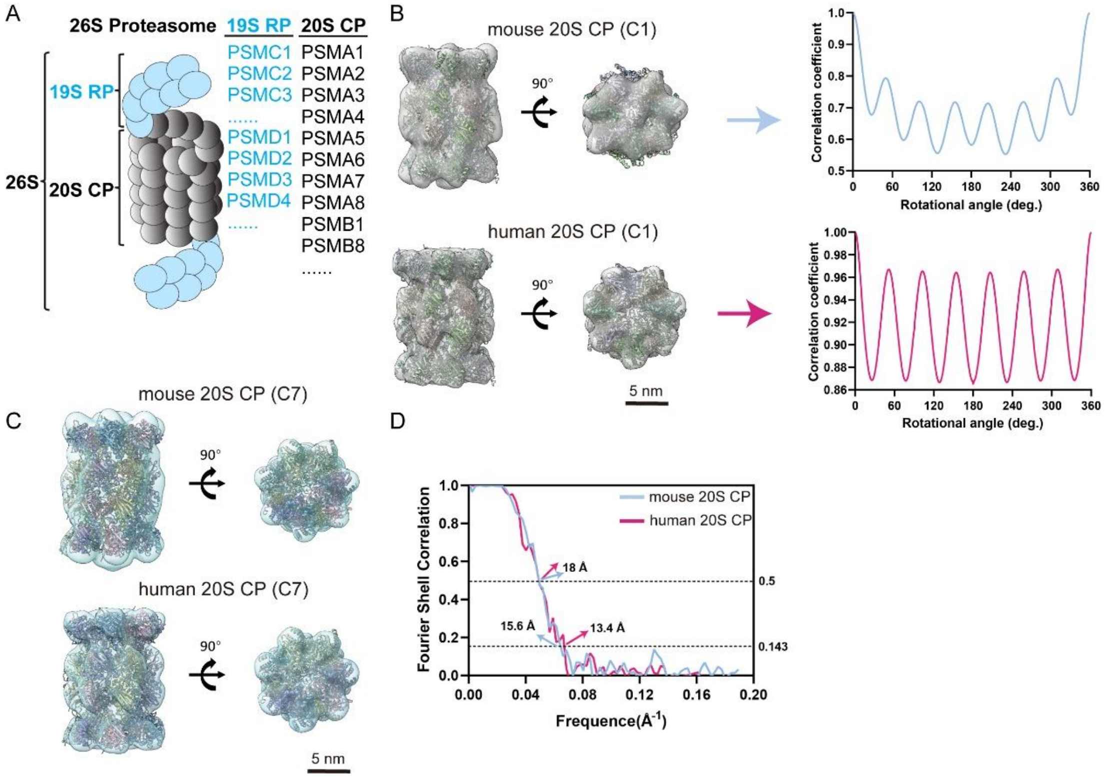


**Figure S1. Structural determination of human and mouse 20S CP**

(A) The 26S Proteasome model. (B) The unsymmetrized average maps fitted well the rat 20S proteasome atomic model (PDB: 6TU3) and showed clear 7-fold symmetry by rotational correlation coefficient analysis (right). (C) Average maps upon application of C7 symmetry (mouse 20S CP, upper; human 20S CP, lower), fitted with rat 20S proteasome atomic model (PDB: 6TU3). (D) Gold-standard Fourier shell correlation curves of the two maps shown in (C) (mouse: 15.6 Å; human: 13.4 Å at 0.143 cutoff. 18 Å for both maps at 0.5 cutoff).

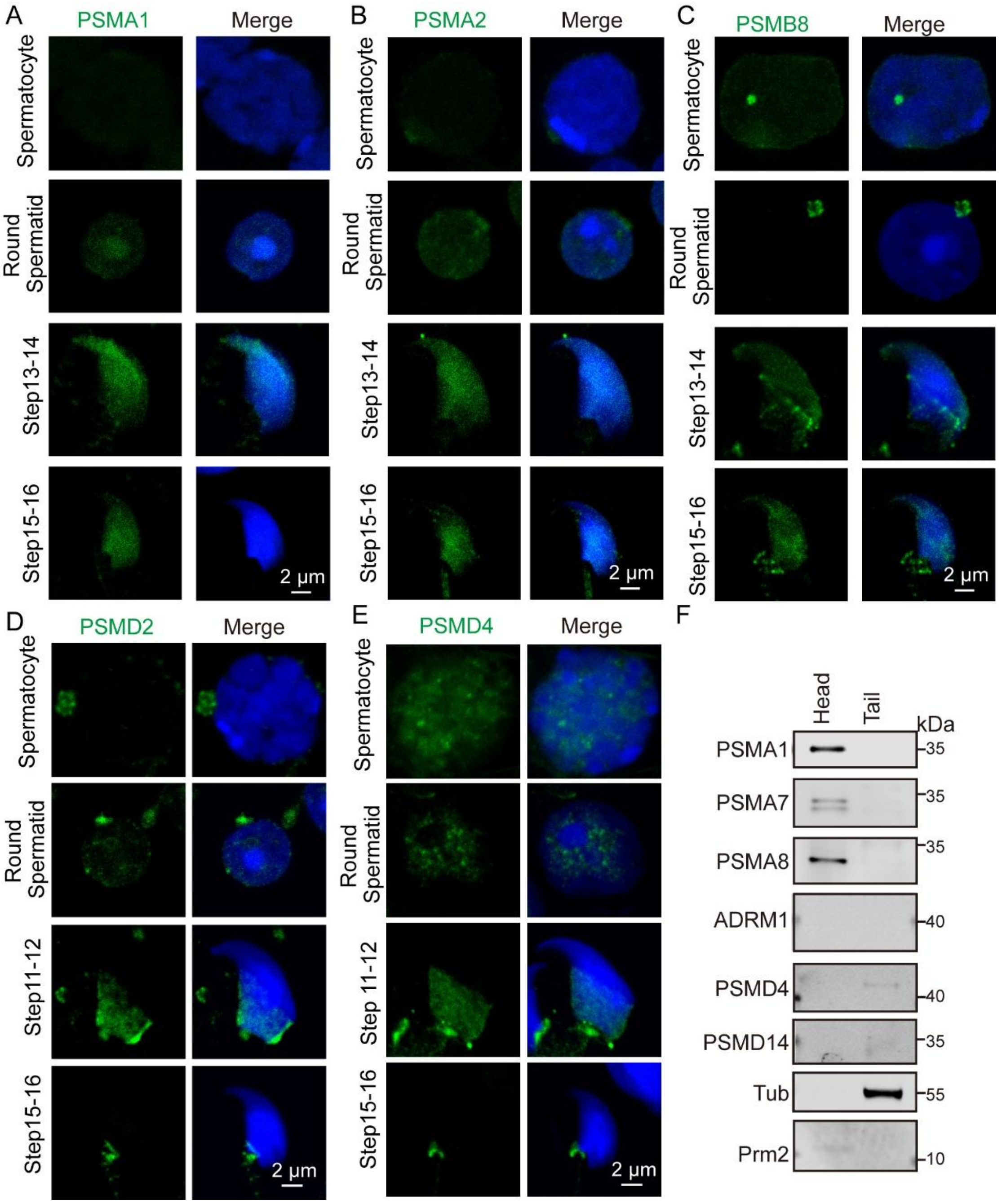


**Figure S2. 20S CP components form foci in the mouse sperm nucleus**

(A-E) Immunofluorescence showing the localization of PSMA1 (A) PSMDA2 (B) PSMB8 (C) PSMD2 (D) and PSMD4 (E) in developing mouse germ cells. (F) The expression of proteasome components in mature mouse sperm.

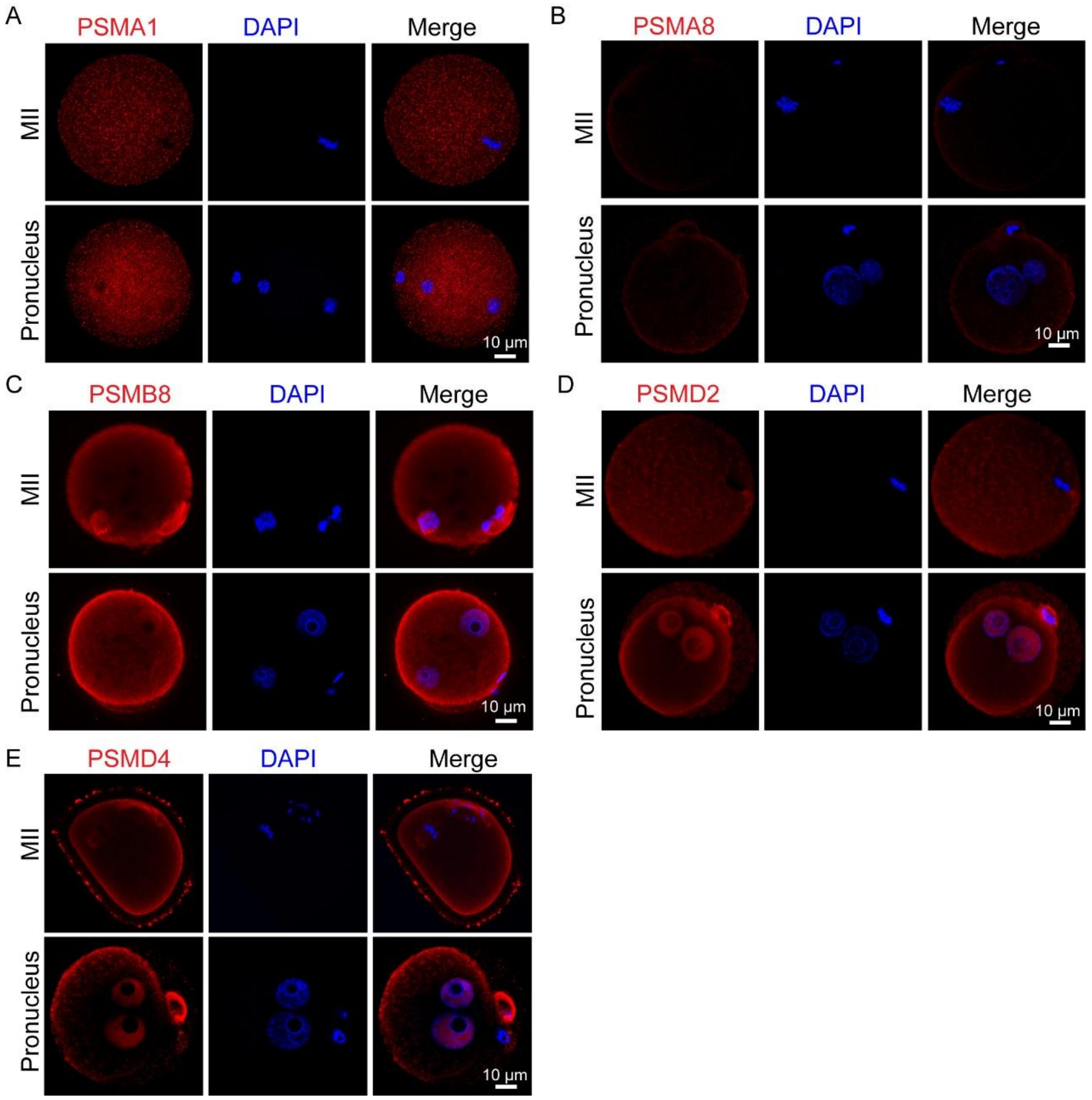


**Figure S3. The expression of proteasome components in mouse oocytes and embryos.**

(A-F) Immunofluorescence showing the expression of PSMA1 (A), PSMA8 (B), PSMB8 (C), PSMD2 (D), and PSMD4 (E) in MII oocytes and pronuclei.

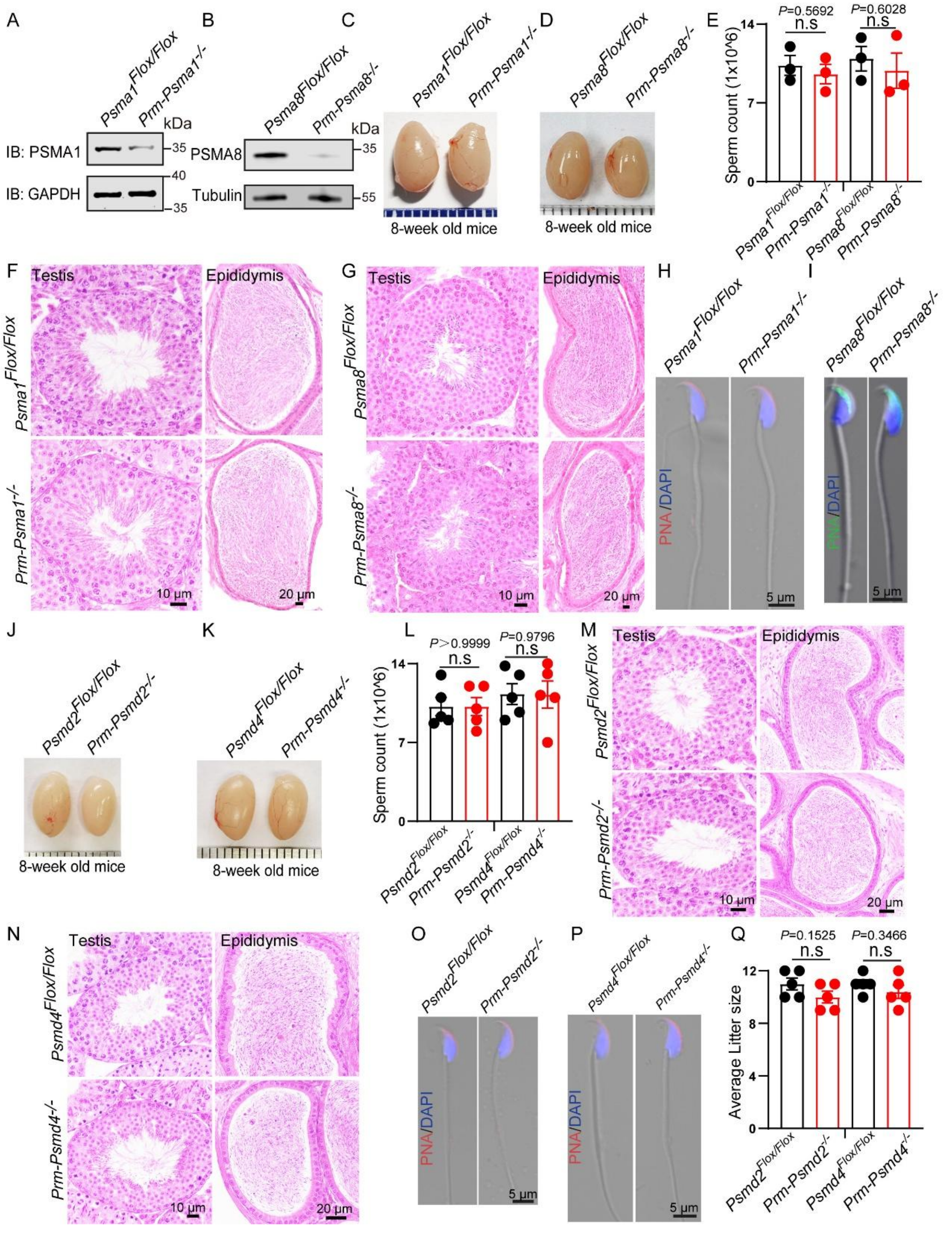


**Figure S4. The spermatogenesis process is normal in proteasome components depletion mice.**

(A and B) PSMA1 or PSMA8 protein levels were dramatically reduced in the sperm of *Prm-Psma1*$^{-/-}$ or *Prm-Psma8*$^{-/-}$mice. Western blots were probed with anti-PSMA1, anti-PSMA8, anti-Tubulin, and anti-GAPDH antibodies. Tubulin and GAPDH served as a loading control. (C and D) Testis size did not differ between 8-week-old *Psma1*$^{Flox/Flox}$ mice and *Prm-Psma1*$^{-/-}$ mice, or between 8-week-old *Psma8*$^{Flox/Flox}$ mice and *Prm-Psma8*$^{-/-}$ mice. (E) Sperm count in the caudal epididymis are similar in each group. *Psma1*$^{Flox/Flox}$, 10.33 ± 0.81 (n = 3 independent experiments); *Prm-Psma1*$^{-/-}$, 9.57 ± 0.87 (n = 3 independent experiments); *Psma8*$^{Flox/Flox}$, 10.93 ± 1.10 (n = 3 independent experiments); *Prm-Psma8*$^{-/-}$, 9.87 ± 1.57 (n = 3 independent experiments). Data are presented as the mean ± SEM. Two-tailed unpaired Student's t test, n.s., non-significant. (F and G) The histomorphology of *Prm-Psma1*$^{-/-}$ and *Prm-Psma8*$^{-/-}$ seminiferous tubules were similar to the control groups as shown by H&E staining of testes in *Psma1*$^{Flox/Flox}$ and *Prm-Psma1*$^{-/-}$, *Psma8*$^{Flox/Flox}$ and *Prm-Psma8*$^{-/-}$ mice. (H and I) Sperm morphology was normal in *Prm-Psma1*$^{-/-}$ and *Prm-Psma8*$^{-/-}$mice. Immunofluorescence staining of PNA in *Psma1*$^{Flox/Flox}$ and *Prm-Psma1*$^{-/-}$, and *Psma8*$^{Flox/Flox}$ and *Prm-Psma8*$^{-/-}$spermatozoa. (J and K) Testis size did not differ between 8-week-old *Psmd2*$^{Flox/Flox}$ and *Prm-Psmd2*$^{-/-}$ mice, or between 8-week-old *Psmd4*$^{Flox/Flox}$ and *Prm-Psmd4*$^{-/-}$ mice. (L) Sperm count in the caudal epididymis was similar in each group. *Psmd2*$^{Flox/Flox}$, 10.20 ± 0.81 (n = 5 independent experiments); *Prm-Psmd2*$^{-/-}$, 10.20 ± 0.80 (n = 5 independent experiments); *Psmd4*$^{Flox/Flox}$, 11.30 ± 0.93 (n = 5 independent experiments); *Prm-Psmd4*$^{-/-}$, 11.26 ± 1.21 (n = 5 independent experiments). Data are presented as the mean ± SEM. Two-tailed unpaired Student's t test, n.s., non-significant. (M and N) The histomorphology of *Prm-Psmd2*$^{-/-}$ and *Prm-Psmd4*$^{-/-}$ seminiferous tubules were similar to the control groups as shown by H&E staining of testes in *Psmd2*$^{Flox/Flox}$ and *Prm-Psmd2*$^{-/-}$, *Psmd4*$^{Flox/Flox}$ and *Prm-Psmd4*$^{-/-}$mice. (O and P) Sperm morphology was normal in *Prm-Psmd2*$^{-/-}$ and *Prm-Psmd4*$^{-/-}$mice. Immunofluorescence staining of PNA (red) in *Psmd2*$^{Flox/Flox}$ and *Prm-Psmd2*$^{-/-}$, *Psmd4*$^{Flox/Flox}$ and *Prm-Psmd4*$^{-/-}$ spermatozoa. (Q) The average litter size of *Psmd2*$^{Flox/Flox}$, 11.00 ± 0.45 (n = 5 independent experiments); *Prm-Psmd2*$^{-/-}$, 10.00 ± 0.45 (n = 5 independent experiments); *Psmd4*$^{Flox/Flox}$, 11.00 ± 0.32 (n = 5 independent experiments); *Prm-Psmd4*$^{-/-}$, 10.40 ± 0.51 (n = 5 independent experiments). Data are presented as the mean ± SEM. Two-tailed unpaired Student's t test, n.s., non-significant.

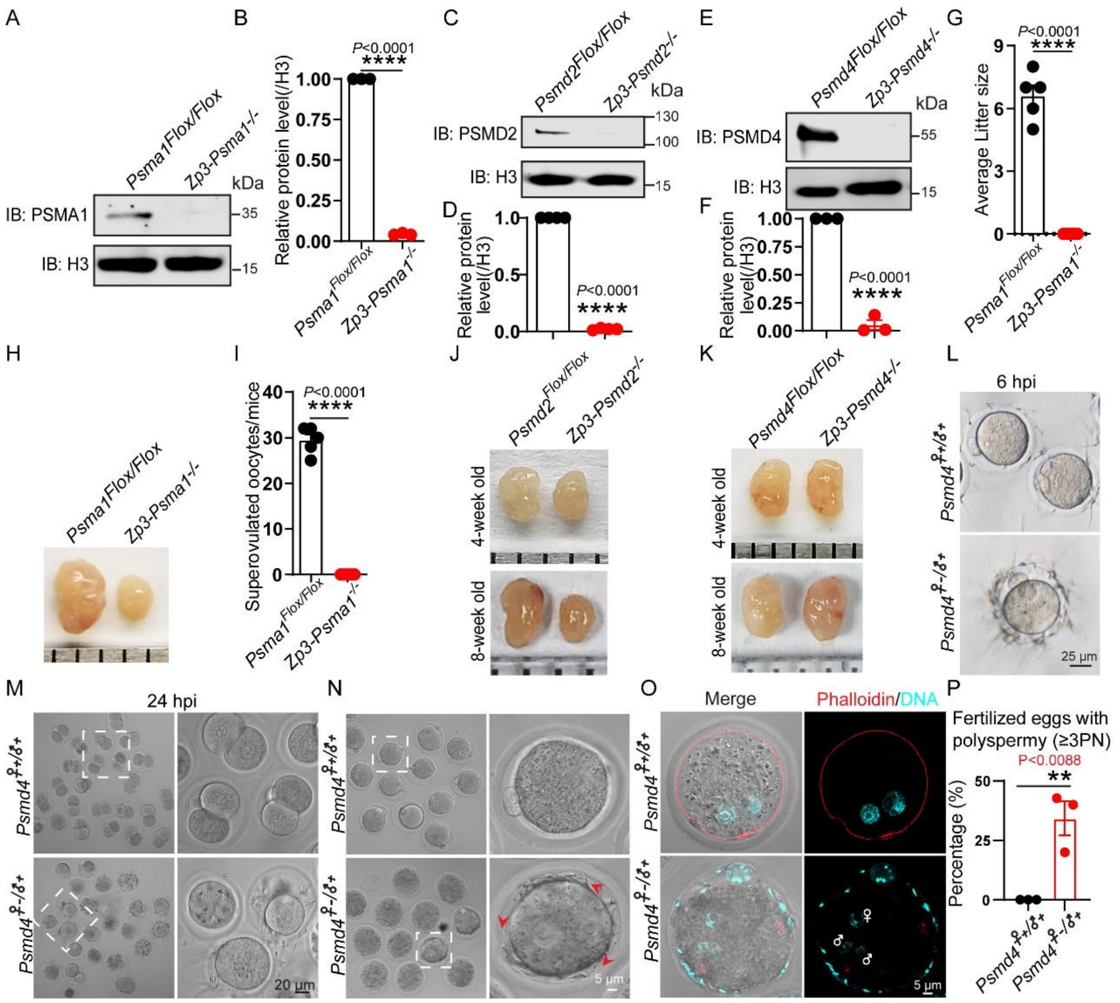


**Figure S5. Depletion of 19S RP components leads to complete infertility in female mice.**

(A) PSMA1 protein levels were dramatically reduced in the oocytes of *Zp3-Psma1*$^{-/-}$ mice. Western blots were probed with anti-PSMA1 and anti-H3 antibodies. H3 served as a loading control. (B) Relative protein levels of PSMA1 in *Psma1*$^{Flox/Flox}$ and *Zp3-Psma1*$^{-/-}$ oocytes. (n = 3 independent experiments). Data are presented as the mean ± SEM. Two-tailed unpaired Student's t test, ****p < 0.0001. (C) PSMD2 protein levels were dramatically reduced in the oocytes of *Zp3-Psmd2*$^{-/-}$mice. Western blots were probed with anti-PSMD2 and anti-H3 antibodies. H3 served as a loading control. (D) Relative protein levels of PSMD2 in (C). (n = 3 independent experiments). Data are presented as the mean ± SEM. Two-tailed unpaired Student's t test, ****p < 0.0001. (E) PSMD4 protein levels were dramatically reduced in the oocytes of *Zp3-Psmd4*$^{-/-}$ mice. Western blots were probed with anti-PSMD4 and anti-H3 antibodies. H3 served as a loading control. (F) Relative protein levels of PSMD4 in (E). (n = 3 independent experiments). Data are presented as the mean ± SEM. Two-tailed unpaired Student's t test, ****p < 0.0001. (G) The average litter size of *Psma1*$^{Flox/Flox}$, 6.60 ± 0.51 (n = 5 independent experiments); *Zp3-Psma1*$^{-/-}$, 0.00 ±

0.00 (n = 5 independent experiments). Data are presented as the mean ± SEM. Two-tailed unpaired Student's t test, ****$p < 0.0001$. (H) Ovaries of 8-week-old *Zp3-Psma1*$^{-/-}$ mice were smaller than those of 8-week-old *Psma1*$^{Flox/Flox}$ mice. (I) Quantitative analysis of oocyte numbers for *Psma1*$^{Flox/Flox}$ and *Zp3-Psma1*$^{-/-}$ mice after superovulation. *Psma1*$^{Flox/Flox}$, 29.40 ± 1.33 (n = 5 independent experiments, total oocytes = 147); *Zp3-Psma1*$^{-/-}$, 0.00 ± 0.00 (n = 5 independent experiments, total oocytes = 0). (J) Ovary size was similar for 4-week-old *Zp3-Psmd2*$^{-/-}$ mice and *Psmd2*$^{Flox/Flox}$ mice. However, ovaries of 8-week-old *Zp3-Psmd2*$^{-/-}$ mice were smaller than those of 8-week-old *Psmd2*$^{Flox/Flox}$ mice. (K) Ovary size was similar for *Zp3- Psmd4*$^{-/-}$ mice and *Psmd4*$^{Flox/Flox}$ mice (4-week-old and 8-week-old). (L) Representative images of embryos in control and maternal *Psmd4*-depleted mice after 6 hours post IVF. (M) Representative images of embryos in control and maternal *Psmd4*-depleted mice after 24 hours post IVF. (N and O) *In vivo* development of embryos from *Psmd4*$^{Flox/Flox}$ and *Zp3-Psmd4*$^{-/-}$ female mice. Embryos from *Psmd4*$^{Flox/Flox}$ and *Zp3-Psmd4*$^{-/-}$female mice were isolated 0.5 days after mating with normal male mice. Representative images of embryos stained with Phalloidin (red) and DAPI (cyan). Arrows indicate extra sperm in the perivitelline space of embryos. Female and male symbols indicate the female and male pronuclei, respectively. (P) Percentage of fertilized oocytes with polyspermy (≥3PN) was quantified in (O). *Psmd4*$^{♀+/♂+}$, 0.00% ± 0.00% (n = 3 independent experiments, total oocytes = 28); *Psmd4*$^{♀-/♂+}$, 34.27% ± 7.18% (n = 3 independent experiments, total oocytes = 27). Data are presented as the mean ± SEM. Two-tailed unpaired Student's t test, **$p < 0.01$

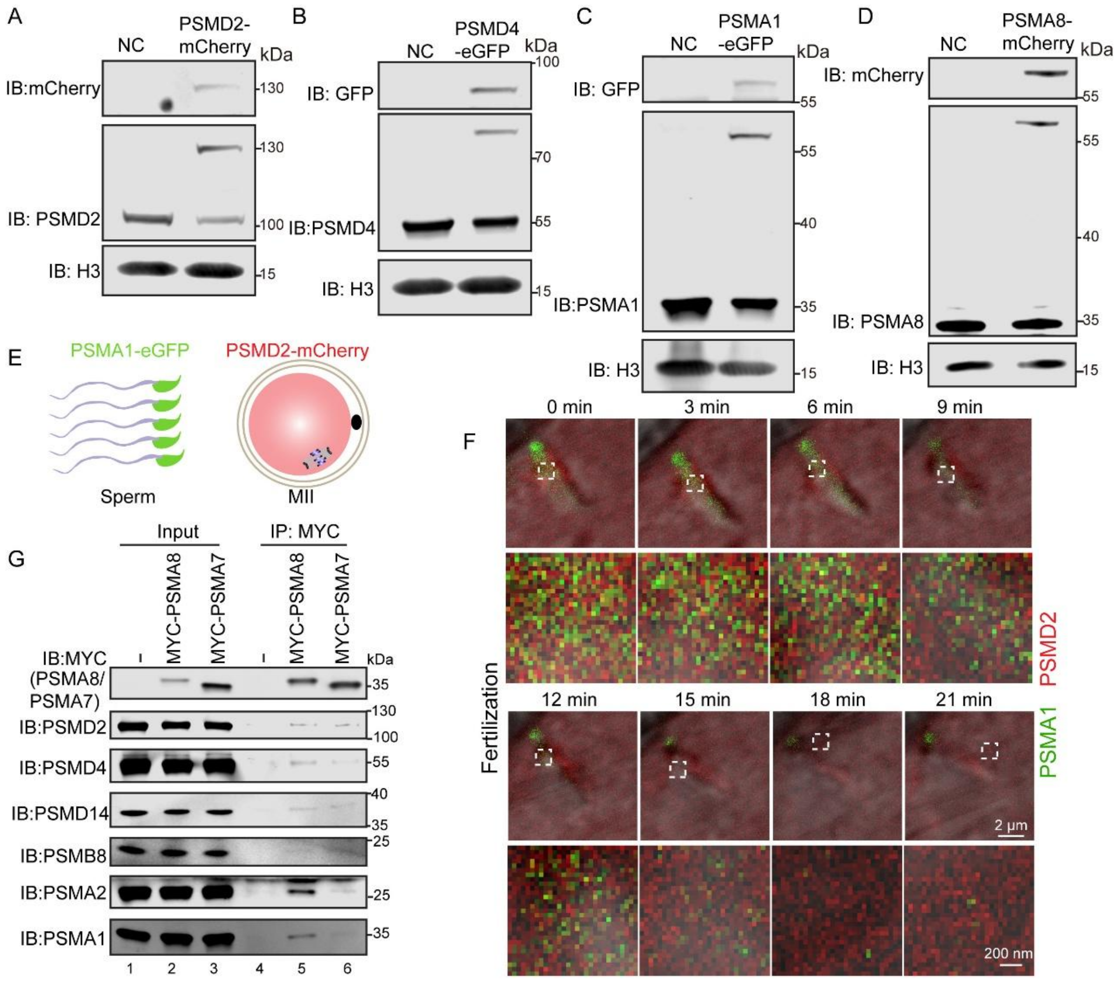


**Figure S6. Testis-specific 20S CP components were integrated with oocyte-derived 19S RP components during fertilization**

(A-D) Western blotting showed that PSMD2-mCherry, PSMD4- eGFP, PSMA1-eGFP, and PSMA8-mCherry were expressed in heterozygous knock-in mice. (E) Schematic of *in vitro* fertilization process by PSMA1-eGFP and PSMD2-mCherry knock-in mice. (F) Live imaging results showed that PSMD2-mCherry rapidly formed foci surrounding PSMA8-eGFP sperm and led to the formation of chimeric proteasomes during fertilization. (G) PSMA8 physically interacts with proteasome components. MYC-PSMA8 or MYC-PSMA7 (or empty vector) were co-transfected into HEK293T cells. 24 hours after transfection, cells were collected for immunoprecipitation with anti-MYC antibody and then visualized using anti-PSMA1, anti-PSMA2, anti-PSMB8, anti-PSMD2, anti-PSMD4, anti-PSMD14, or anti-MYC antibodies.

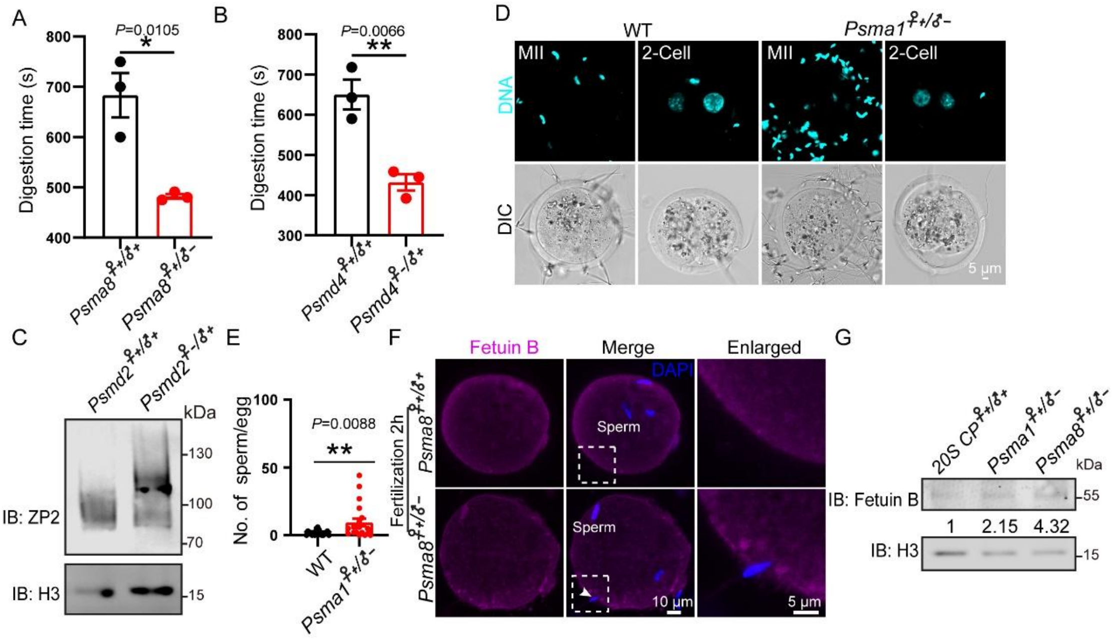


**Figure S7. Depletion of proteasome components prevented zona pellucida hardening**

(A) Zona pellucida digestion time in *Psma8*$^{♀+/♂+}$ (n = 3 independent experiments, total embryos = 60) and *Psma8*$^{♀+/♂-}$ (n = 3 independent experiments, total embryos = 60) groups. Data are presented as the mean ± SEM. Two-tailed unpaired Student's t test, $^{*}P < 0.05$. (B) Zona pellucida digestion time in *Psmd4*$^{♀+/♂+}$ (n = 3 independent experiments, total embryos = 60) and *Psmd4*$^{♀-/♂+}$ (n = 3 independent experiments, total embryos = 60) groups. Data are presented as the mean ± SEM. Two-tailed unpaired Student's t test, $^{**}P < 0.01$. (C) ZP2 remains uncleaved in maternal PSMD2-depleted embryo. Western blots were probed with anti-ZP2, and anti-H3 antibodies. H3 served as a loading control. (D) Representative images of sperm binding to the zona pellucida surrounding WT and paternal PSMA1-depleted oocytes and embryos. MII oocytes and two-cell embryos from WT mice were incubated with both WT and PSMA1-depleted capacitated sperm for 1 hour. The oocytes and embryos were then stained with DAPI (cyan). (E) The number of sperm binding to the surface of zona pellucida surrounding oocytes from WT and paternal PSMA1-depleted groups was counted. WT, 1.65 ± 0.40 (n = 19); *Psma1*$^{♀+/♂-}$, 9.50 ± 2.82 (n = 19). Data are presented as the mean ± SEM. Two-tailed unpaired Student's t test, $^{**}p < 0.01$. (F) Representative images showing Fetuin B localization in *Psma8*$^{♀+/♂+}$and *Psma8*$^{♀+/-}$ embryos. Arrows indicate extra sperm in embryos. (G) The Fetuin B protein levels in *Psma1*$^{♀+/-}$ and *Psma8*$^{♀+/-}$ embryos. Western blots were probed with anti-Fetuin B and anti-H3 antibodies. H3 served as a loading control.

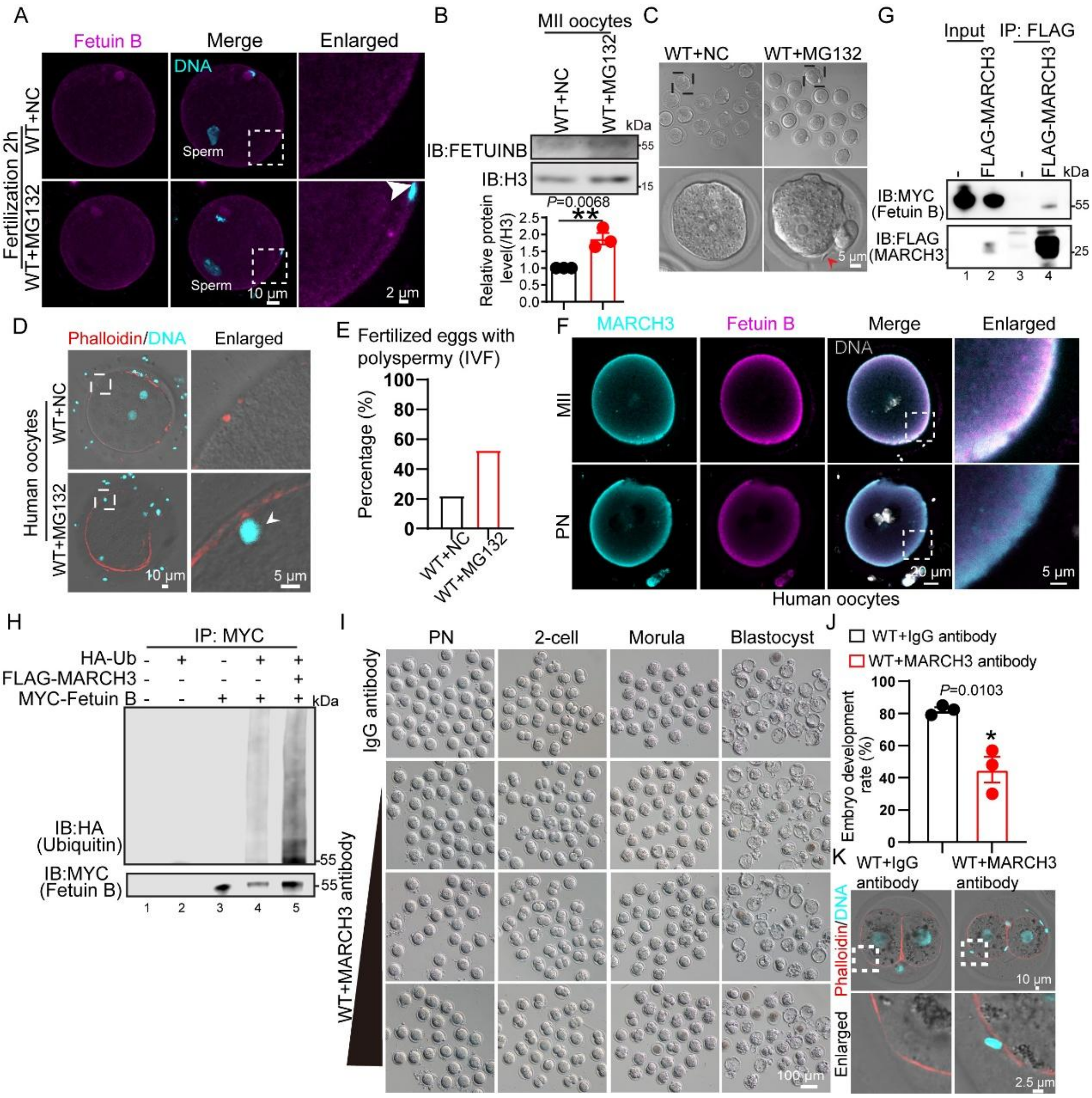


**Figure S8. Inhibiting MARCH3 results in the presence of extra sperm in the perivitelline space of embryos**

(A) Representative images showing the localization of Fetuin B in control and MG132-treated mouse embryos. Arrows indicate extra sperm in embryos. (B) Fetuin B protein levels in MG132-treated oocytes were significantly accumulated compared to those in the control group. Western blots were probed with anti-Fetuin B and anti-H3 antibodies. H3 served as a loading control. (n = 3 independent experiments). Data are presented as the mean ± SEM. Two-tailed unpaired Student's t test, $**p < 0.01$. (C) Representative images of embryos in DMSO and MG132-treated groups after 6 hours post IVF. Arrows indicate extra sperm in the perivitelline space. (D) Representative images of human embryos stained with Phalloidin (red) and DAPI (cyan) in DMSO and MG132 treated group after 8 hours post IVF. Arrows indicate extra human sperm inside the plasma membrane. (E) The percentage of fertilized eggs with extra sperm in the perivitelline space in (D). WT+NC (total human oocytes = 9); WT+MG132 (total oocytes = 8). (F) Representative images showing the localization of Fetuin B and MARCH3 in human oocytes and embryos at the indicated developmental stages. (G) Fetuin B physically interacts with MARCH3. MYC-Fetuin B and MARCH3-FLAG (or empty vector) were co-transfected into HEK293T cells. 24 hours after transfection, cells were collected for immunoprecipitation with an anti-FLAG antibody, followed by visualization using anti-FLAG and anti-MYC antibodies. (H) MARCH3 promotes the polyubiquitination of Fetuin B. HEK293T cells were transfected with MYC-Fetuin B, MARCH3-FLAG, and HA-Ub. 24 hours after transfection, cells were collected for immunoprecipitation with an anti-MYC antibody, followed by visualization with anti-HA and anti-MYC antibodies. (I) Distribution of stages of embryos cultured *in vitro* from pronuclear until blastocyst in control IgG and MARCH3 antibody treatment groups after IVF. (J) Percentage of blastocyst development in control IgG and MARCH3 antibody treatment groups. WT+ IgG antibody, 82.13% ± 1.74% (n = 3 independent experiments, total oocytes = 99); WT+ MARCH3 antibody, 45.00% ± 7.94% (n = 3 independent experiments, total oocytes = 135). Data are presented as the mean ± SEM. Two-tailed unpaired Student's t test, *$p < 0.05$. (K) Representative images of mouse embryos stained with Phalloidin (red) and DAPI (cyan) in control IgG antibody and MARCH3 antibody treatment groups after 24 hours post IVF.

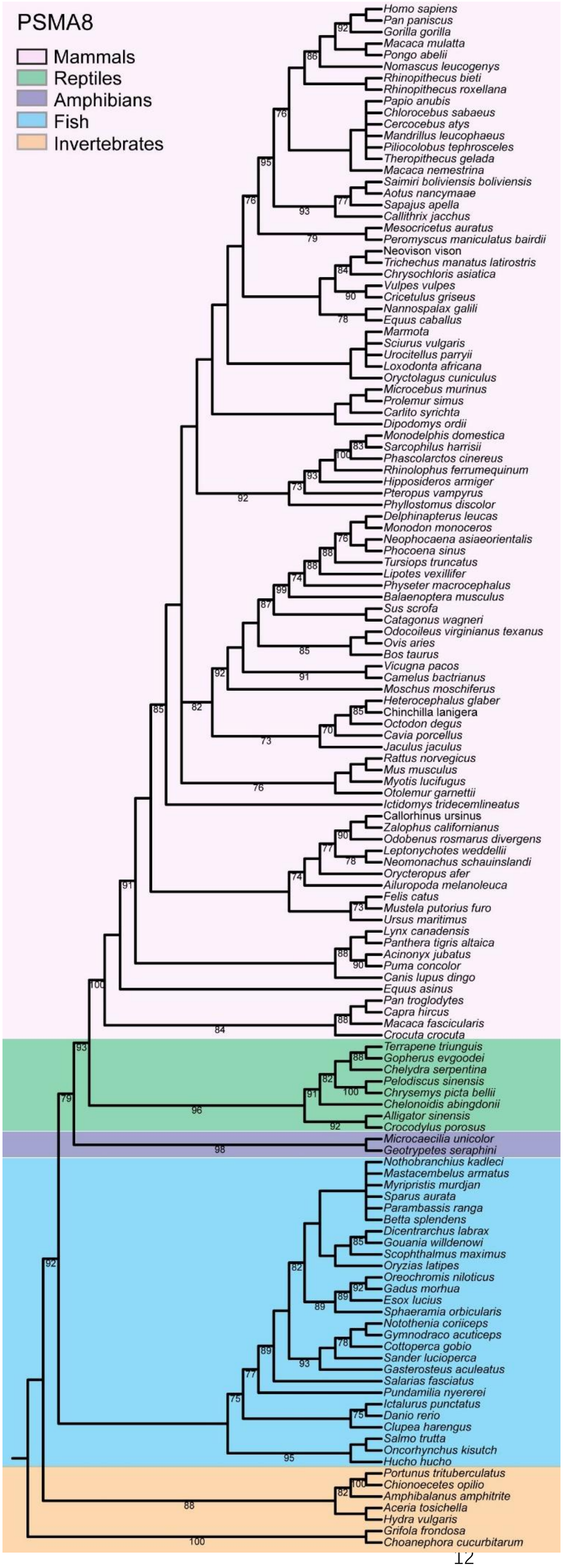
PSMA8
Mammals
Reptiles
Amphibians
Fish
Invertebrates
Homo sapiens
Pan paniscus
Gorilla gorilla
Macaca mulatta
Pongo abelii
Nomascus leucogenys
Rhinopithecus bieti
Rhinopithecus roxellana
Papio anubis
Chlorocebus sabaeus
Cercocebus atys
Mandrillus leucophaeus
Piliocolobus tephrosceles
Theropithecus gelada
Macaca nemestrina
Saimiri boliviensis boliviensis
Aotus nancymaae
Sapajus apella
Callithrix jacchus
Mesocricetus auratus
Peromyscus maniculatus bairdii
Neovison vison
Trichechus manatus latirostris
Chrysochloris asiatica
Vulpes vulpes
Cricetulus griseus
Nannospalax galili
Equus caballus
Marmota
Sciurus vulgaris
Urocitellus parryii
Loxodonta africana
Oryctolagus cuniculus
Microcebus murinus
Prolemur simus
Carlito syrichta
Dipodomys ordii
Monodelphis domestica
Sarcophilus harrisii
Phascolarctos cinereus
Rhinolophus ferrumequinum
Hipposideros armiger
Pteropus vampyrus
Phyllostomus discolor
Delphinapterus leucas
Monodon monoceros
Neophocaena asiaeorientalis
Phocoena sinus
Tursiops truncatus
Lipotes vexillifer
Physeter macrocephalus
Balaenoptera musculus
Sus scrofa
Catagonus wagneri
Odocoileus virginianus texanus
Ovis aries
Bos taurus
Vicugna pacos
Camelus bactrianus
Moschus moschiferus
Heterocephalus glaber
Chinchilla lanigera
Octodon degus
Cavia porcellus
Jaculus jaculus
Rattus norvegicus
Mus musculus
Myotis lucifugus
Otolemur garnettii
Ictidomys tridecemlineatus
Callorhinus ursinus
Zalophus californianus
Odobenus rosmarus divergens
Leptonychotes weddellii
Neomonachus schauinslandi
Orycteropus afer
Ailuropoda melanoleuca
Felis catus
Mustela putorius furo
Ursus maritimus
Lynx canadensis
Panthera tigris altaica
Acinonyx jubatus
Puma concolor
Canis lupus dingo
Equus asinus
Pan troglodytes
Capra hircus
Macaca fascicularis
Crocuta crocuta
Terrapene triunguis
Gopherus evgoodei
Chelydra serpentina
Pelodiscus sinensis
Chrysemys picta bellii
Chelonoidis abingdonii
Alligator sinensis
Crocodylus porosus
Microcaecilia unicolor
Geotrypetes seraphini
Nothobranchius kadleci
Mastacembelus armatus
Myripristis murdjan
Sparus aurata
Parambassis ranga
Betta splendens
Dicentrarchus labrax
Gouania willdenowi
Scophthalmus maximus
Oryzias latipes
Oreochromis niloticus
Gadus morhua
Esox lucius
Sphaeramia orbicularis
Notothenia coriiceps
Gymnodraco acuticeps
Cottoperca gobio
Sander lucioperca
Gasterosteus aculeatus
Salarias fasciatus
Pundamilia nyererei
Ictalurus punctatus
Danio rerio
Clupea harengus
Salmo trutta
Oncorhynchus kisutch
Hucho hucho
Portunus trituberculatus
Chionoecetes opilio
Amphibalanus amphitrite
Aceria tosichella
Hydra vulgaris
Grifola frondosa
Choanephora cucurbitarum

**Figure S9. PSMA8 is evolutionarily conserved in mammals**

The phylogenetic tree in this figure represents a comparative analysis of PSMA8 sequences in representative organisms obtained from the Universal Protein Resource (UniProt) Knowledgebase.

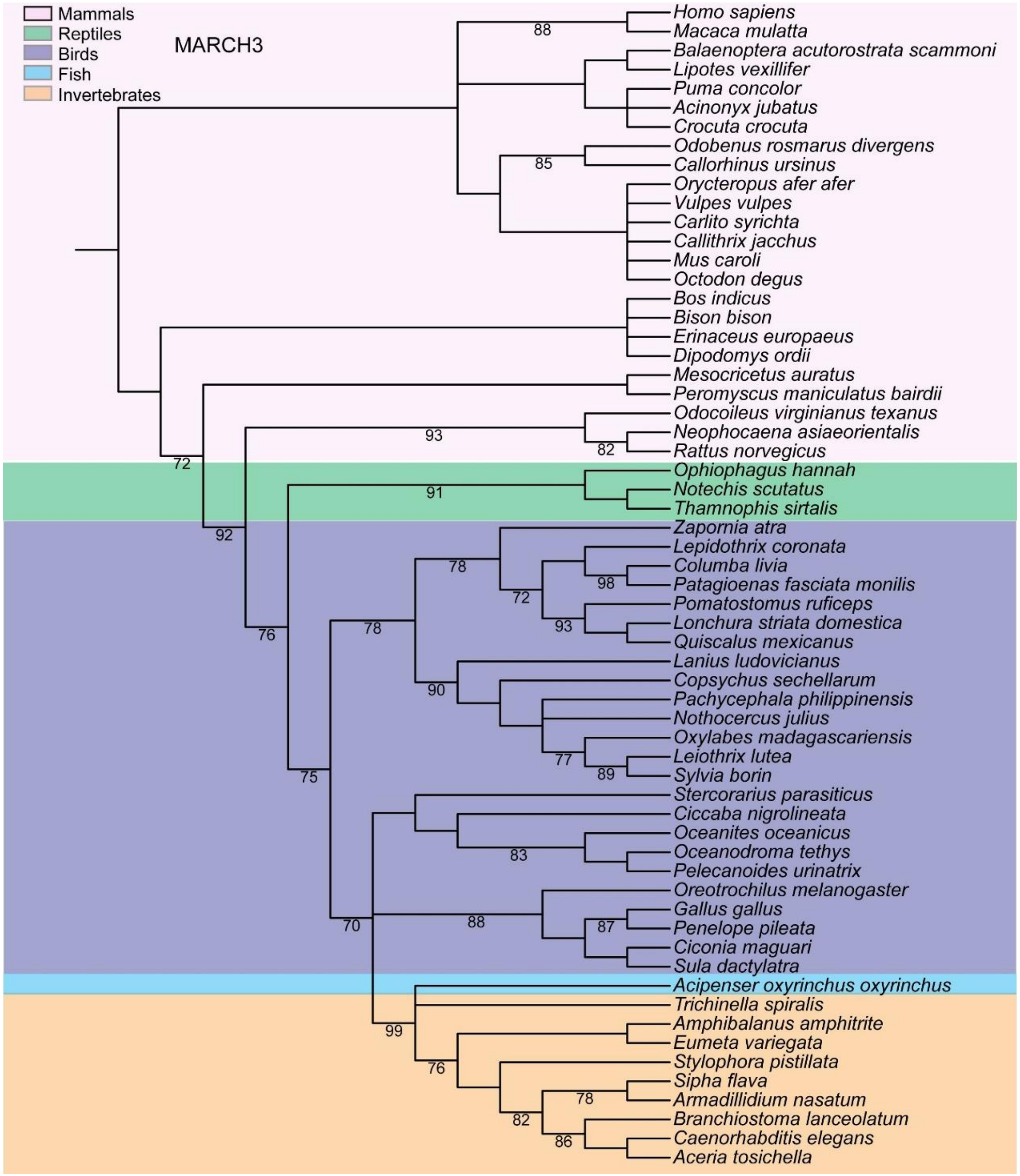


**Figure S10. MARCH3 is evolutionarily conserved from worm to mammals**

The phylogenetic tree in this figure represents a comparative analysis of MARCH3 sequences in representative organisms obtained from the Universal Protein Resource (UniProt) Knowledgebase.
